# Diversity and evolution of a phase-variable multi-locus antigen in *Neisseria gonorrhoeae*

**DOI:** 10.64898/2026.02.02.703239

**Authors:** QinQin Yu, Tatum D. Mortimer, Sofia O.P. Blomqvist, Bailey Bowcutt, David Helekal, Samantha G. Palace, Yonatan H. Grad

## Abstract

*Neisseria gonorrhoeae* is a sexually transmitted bacterial pathogen that deploys multiple mechanisms to evade the immune system, including rapid variation in surface antigens. One of the most abundant and diverse antigens is the Opacity (Opa) protein, a surface protein that mediates gonococcal attachment to host receptors. Studies of Opa diversity and evolution have been limited by the inability of short-read sequencing to resolve the multiple copies of *opa* in each genome, preventing a comprehensive understanding of antigenic variation for vaccine design and immunology studies. We assembled a dataset of 219 complete genomes from diverse clinical isolates using long-read sequencing and developed bioinformatics and phylogenetics tools to assess *opa* variation quantitatively. Each genome had on average 7 distinct *opa* alleles at 9 to 12 *opa* loci, and almost all isolates had at least one pair of identical or near-identical *opa* genes. Fewer *opa* genes were in frame, and thus inferred to be expressed, than expected due to chance. While genomic distance between isolates correlated with overall *opa* allele sequence similarity, *opa* genes were on average 74 times more diverse than the rest of the genome. One *opa* locus evolved more rapidly than the other loci. There was little evidence that interspecies recombination contributed to *N. gonorrhoeae opa* diversity. Our findings reveal a continuously evolving *opa* repertoire that leads to diverse *opa* alleles even in closely related strains and indicate that there are likely unknown biological factors modulating *opa* expression.

**Author Summary:** The rising levels of antibiotic resistance in *Neisseria gonorrhoeae* make controlling the spread of this sexually transmitted pathogen a public health priority. *N. gonorrhoeae* rapidly varies surface proteins to evade recognition by the human adaptive immune system. Understanding how these proteins evolve may help us design better vaccines and control measures for curbing the spread of gonorrhea. One of the most abundant surface proteins is the Opacity protein (Opa), which helps *N. gonorrhoeae* bind to host cells upon colonization. Research efforts to understand the evolution of Opa have been limited because it is encoded by multiple genes in the genome that are not resolved by short-read sequencing technologies. Here, we resolved the genes that encode Opa using a dataset of 132 publicly available complete genomes and 87 genomes that we completed using long-read sequencing of diverse clinical isolates. We found that Opa evolves rapidly to generate different versions of the protein in the same isolate, but very few of these protein versions appear expressed. We also found evidence that there may be other, uncharacterized mechanisms that control how these proteins evolve over longer timescales.

## Introduction

In 2020, the World Health Organization (WHO) estimated there were more than 82 million cases worldwide of gonorrhea, the sexually transmitted infection caused by the obligate human pathogen *Neisseria gonorrhoeae*. Gonorrhea is a WHO priority pathogen due to increasing levels of antibiotic resistance and no highly-effective vaccine [1]. Infection does not appear to confer protection, such that individuals can be reinfected [2], even by the same strain [3], reflecting *N. gonorrhoeae*’s escape from the host immune system [4,5].

One of the ways *N. gonorrhoeae* achieves this immune evasion is through rapidly varying its antigens [6,7]. A major surface protein that undergoes this variation is the Opacity protein (Opa) [8–11]. Opa proteins mediate gonococcal attachment to host receptors during colonization [12–17] and may be involved in suppression of the host immune response, though the mechanisms are as yet unclear [15,18]. These proteins are encoded by multiple highly diverse *opa* genes in separate loci in each *N. gonorrhoeae* genome [10,19]. A series of pentanucleotide repeats in the 5’ end of the coding sequence determines whether each of these genes is in frame and fully translated, and changes in the repeat copy number, likely from slip-strand repair, result in phase variation [20].

Studies of *opa* variation and evolution have focused on small numbers of isolates or closely related isolates [9,19,21]. These studies showed that *opa* genes have high sequence diversity within and between isolates [9], recombination within isolates plays a key role in generating new sequence diversity [11,21], and gene conversion events can lead to two identical or near-identical *opa* alleles [20–22]. Bilek et al. [21] observed that one *opa* gene varied more in closely related isolates than the other *opa* loci. There is also evidence that *opa* loss occasionally occurs [21] and few *opa* are phase on in each isolate [20]. Phylogenetic analysis of *opa* from *N. gonorrhoeae, N. meningitidis,* and the commensals *N. flava* and *N. sicca* showed that *opa* clustered by species [9].

However, due to the small numbers of isolates in these studies, we lack generalizable quantitative patterns and thus have a limited understanding of *N. gonorrhoeae opa* sequence variation and evolution. Several obstacles have prevented studies of *opa* diversity in larger numbers of isolates. Polymerase chain reaction can amplify the *opa* regions [19,21], but this approach is challenging to scale up due to unknown primer coverage of the *opa* repertoire. *opa* sequences cannot be resolved using short-read whole-genome sequencing platforms, such as Illumina, as the multiple and highly similar *opa* loci are problematic for mapping and genome assembly. Furthermore, the high diversity of *opa* has made it challenging to analyze sequences using standard bioinformatics, phylogenetics, and phylodynamic approaches, in that methods are designed primarily for single copy genes.

Long-read sequencing enables us to overcome these challenges. We combined publicly available complete genomes with diverse genomes that we sequenced using the Oxford Nanopore long-read platform. We developed bioinformatic and phylogenetic approaches for defining the diversity and evolution of highly variable multilocus antigens and applied them to the largest set of *opa* sequences to date to investigate *opa* variation. Finally, we developed a sequence-based clustering approach for *opa* alleles in the effort towards establishing a standardized nomenclature to facilitate comparison of *opa* alleles in future studies.

## Methods

### Dataset of high-quality complete genomes

#### Publicly available complete genomes

We searched for “*Neisseria gonorrhoeae*” in the National Center for Biotechnology Information Genome Datasets (https://www.ncbi.nlm.nih.gov/datasets/genome/) and downloaded all genomes with an assembly level of “chromosome” or “complete” on July 31, 2025. We determined the sequencing technology and assembly method from the associated publications (where available) and associated metadata available on NCBI. In the rare cases of discrepancies between the publication and NCBI, we used the metadata reported in the publication.

#### Nanopore sequencing and genome assembly

*N. gonorrhoeae* isolates were streaked from frozen stocks onto GCB agar (Difco) with Kellogg’s supplement [23] and grown at 37° C with 5% CO_2_ overnight. Genomic DNA was extracted using the Invitrogen PureLink Genomic DNA Mini Kit. We prepared the library using the Oxford Nanopore Native Barcoding Kit 24 V14 (SQK-NBD114.24) and performed Nanopore sequencing using a MinION Mk1B device. Basecalling was performed using Dorado v0.8.2 [24] superaccuracy basecalling. Reads were demultiplexed using Dorado v0.8.2 demux and filtered using Filtlong v0.2.1 (https://github.com/rrwick/Filtlong) to keep the 90% highest quality read bases and only reads longer than 1kbp. Genomes were assembled using Autocycler v0.2.1 [25] with 4 read subsets at 25x minimum depth and using the assemblers: Canu v2.2 [26], Flye v2.9.5 [27], miniasm v0.3 [28], NECAT v0.0.1 [29], NextDenovo v2.5.2 [30], and Raven v1.8.3 [31]. Genomes that were fully resolved (one complete genome plus optional plasmids) and had a length of at least 2 Mbp were kept, as we expect the length of the *N. gonorrhoeae* genome to be 2.2 Mbp.

### Identifying *opa* genes in complete genomes

Genomes were rotated using the rotate program v1.0 [32] to match the first 90 bases of *dnaA* from FA1090 (NC_002946.2), allowing for up to 5 mismatches. We wrote a custom script to identify *opa* genes that searched for the tandem pentanucleotide repeats (CTCTT) and a unique conserved sequence near the stop codon [19,21]. Details of the algorithm are presented in the Supplementary Information. The N-terminus of the mature protein was found by searching for the sequence GCAAGTGA [20], allowing at most 2 substitutions. Only sequences that have an identified start codon, stop codon, and N-terminus are determined to be intact *opa* genes. Intact *opa* genes were given a number starting from 1 in the order that they appeared in the rotated genome (i.e., FA1090 *opa1* is the first *opa* in FA1090 after *dnaA*). The *opa* sequence was determined to be in frame if the number of nucleotides between the start codon and the N-terminus was a multiple of 3.

### Comparison to other sequence search methods

To confirm that we had identified all the *opa* genes in the genomes, we compared our method to three additional approaches:

1. We performed a search of the conserved region of the FA1090 *opa1* sequence between the semivariable and hypervariable 1 regions using BLAST v2.14.1. We set the maximum number of high-scoring segment pairs (local alignments) to keep as 15 for a single query-subject pair and kept all other parameters at the default values. We matched the hits from the BLAST results that were also found by our search algorithm by merging the nearest start positions of the hits from the two search algorithms if they were within 1200 nucleotides. BLAST hits that were not merged were identified as putative additional hits found by BLAST not found by our search algorithm and we manually inspected these sequences.
2. We annotated the genomes using Prokka v1.14.6 [33]. We set the genus to “Neisseria” and species to “gonorrhoeae” and kept all other parameters at their default values. We noticed that all the *opa* genes identified with our method had the gene name *piiC*_*, where * was a number. We then checked if there were any additional genes annotated as *piiC_*\* that were not identified by our method.
3. We used Roary v3.13.0 [34] to define the pan-genome and cluster genes. We set the minimum percentage identity for the BLASTp program within Roary to be 90% and kept all other parameters at their default values. We looked at gene clusters that included the *opa* genes identified by our method to assess for additional genes in those clusters not identified by our method.

### Identification of truncated opa

To identify truncated *opa* that were missing the start and/or stop codon, we extracted the sequences, including upstream and downstream sequences, of the hits found by BLAST that were not found by our search algorithm and aligned the sequences with the FA1090 *opa* sequences using MAFFT v7.520 [35]. We visualized the alignments using JalView v2.11.5.1 [36] to identify if each hit was a truncated *opa* compared to the FA1090 *opa* sequences.

### Genomic rearrangements

Genomic rearrangements were detected using progressiveMauve v2.4.0 [37] by comparing each complete genome (query) with the complete FA1090 genome (reference). We rearranged the *opa* position in the query genome relative to the reference genome by reordering and flipping regions of the query genome that fell into locally collinear blocks compared to the reference. Regions of the query genome that did not fall into any locally collinear block with the reference genome were not altered.

### Whole genome phylogenies

#### Generating reference-mapped pseudogenomes

To generate reference-mapped pseudogenomes, we used short reads generated by, or simulated to be generated by, Illumina sequencing. Sequencing reads from all publicly available *N. gonorrhoeae* datasets were downloaded from the European Nucleotide Archive (**Table S3** and https://github.com/qinqin-yu/gonococcus_opa_diversity/blob/main/opa_diversity_snakemake/input_data/whole_genome_ metadata/global_isolates_metadata_met_qc.csv). For the publicly available complete genomes, we used ART v2016.06.05 [38] to simulate paired-end short reads from the MiSeq v3 sequencing system with a read length of 250 bp, 80x coverage, mean DNA fragment size of 600 and standard deviation of 100. Reads were mapped to the NCCP11945 (NC_011035.1) reference genome using BWA-MEM v0.7.17 [39]. Duplicate reads were marked with Picard v3.0.0 (https://broadinstitute.github.io/picard/), and reads were sorted with SAMtools v1.17 [40]. The quality of the mapped reads was assessed using Qualimap’s bamqc v2.2.1 [41]. We used Pilon v1.24 [42] to call variants (minimum mapping quality 20 and minimum coverage of 10). To create pseudogenomes, we replaced the reference allele with high quality variant calls (at least 90% of reads supporting the allele). Positions called as deletions by Pilon were replaced with a gap character, and positions with low coverage or an indeterminate allele were replaced with an N.

#### Selecting representative global genomes

To select representative global genomes from the publicly available *N. gonorrhoeae* isolates, we included isolates with genomes meeting the following quality control filters: at least 80% of reads mapped to the *N. gonorrhoeae* reference genome, the coverage of reads mapped to the *N. gonorrhoeae* reference genome was >40X, fewer than 12% of the sites in the *N. gonorrhoeae* reference genome were unable to be confidently called using our assembly pipeline, and the *de novo* assembly length was 1.75 Mbp-2.5 Mbp. This yielded 21,653 genomes which were then clustered using PopPUNK v2.6.0 [43]. The isolate in each PopPUNK cluster with the fewest contigs in its *de novo* assembly was chosen as the representative isolate from that cluster, resulting in 737 representative genomes (**Table S4**).

#### Recombination-masked phylogenies

Recombination-masked phylogenetic trees were created using the reference-mapped pseudogenomes in Gubbins v3.3.4 [44] using the GTR substitution model, the RAxML Next Generation tree builder [45], a maximum of 20 iterations for the phylogeny of complete genomes, and a maximum of 5 iterations for the phylogeny of representative and complete genomes.

### Quantifying *opa* diversity

#### Comparison of within- and between-isolate opa diversity

To compare within- and between-isolate *opa* diversity, we first translated the *opa* sequences downstream of the coding repeats. We excluded all sequences with premature stop codons due to frameshift mutations downstream of the coding repeats or nonsense mutations. We then aligned the amino acid sequences using MAFFT v7.520 [35] with the default parameters. We then calculated pairwise distances in the alignment for all pairs of *opa* sequences from different isolates (between-isolate comparison) and all pairs of *opa* sequences from the same isolate (within-isolate comparison).

#### Similar opa within the same isolate

To identify clusters of similar *opa* sequences from the same strain, we used the NetworkX Python package (https://github.com/networkx/networkx) to create graphs where the nodes are the *opa* sequences and the edges connect *opa* sequences with less than 5% difference in amino acid sequence alignment. The clusters of similar *opa* sequences were identified as the connected components in the graph. The percentage of isolates with at least one pair of similar *opa* was determined by finding the isolates that contained at least one *opa* sequence with less than 5% difference in amino acid sequence alignment with another *opa* sequence in the same strain.

#### Comparison of opa sequence distance and genomic sequence distance

We calculated the *opa* sequence distance and the genomic distance between pairs of genomes. To facilitate quantitative comparisons between pairs of genomes, only pairs of genomes with the same number of *opa* genes were kept. The *opa* sequence distance was calculated using the alignment generated above and pairing the *opa* sequences between the two genomes that had the fewest number of segregating sites using a greedy algorithm (i.e., find the two closest *opa* sequences, then find two next closest *opa* sequences, etc. If there are identical matches, then one is chosen at random to be matched.). The total *opa* sequence distance was calculated by summing the distances between each pair of *opa* sequences and dividing by the number of *opa* genes (where a distance of 0 indicates the *opa* repertoire in the two genomes are identical and a distance of 1 indicates that all sites in all *opa* genes in the two genomes are different).

The pairwise genomic sequence distance was calculated as the number of SNPs between pairs of pseudogenomes divided by the length of the pseudogenome using pairsnp v0.3.1 (https://github.com/gtonkinhill/pairsnp). Missing sites (indicated by N) were excluded in the calculation of the SNP distance. Sites that were missing in some isolates but present in the two isolates being compared were included.

### *Other* Neisseria *species* opa

#### Identifying opa genes

We downloaded complete genomes (“chromosome” or “complete” assembly levels) from NCBI RefSeq on February 13, 2025 using the search term “Neisseria” and filtered out all *N. gonorrhoeae* genomes (**Table S5**). Genomes were rotated using the rotate program v1.0 [32] to match the first 90 bases of *dnaA* from FA1090 (NC_002946.2), allowing for up to 5 mismatches. If this sequence was not found, then the unrotated genome was used for downstream analysis. This occurred in 60/198 (30%) genomes. We identified *opa* genes using the custom script that we used for *N. gonorrhoeae*, which looks for the coding repeats and the unique conserved sequence near the stop codon. To look for additional *opa* that were missed by our script, we BLASTed the conserved regions of *N. gonorrhoeae* FA1090 *opa1*.

#### Assessing sequence relatedness

We calculated the k-mer distances using MASH v2.3 [46] with a k-mer size of 9 and used the distances as input to create a neighbor joining tree using rapidNJ v2.3.2 [47].

### Clustering of variable region sequences

#### Definition of semivariable, hypervariable 1, and hypervariable 2 region sequences

We aligned the nucleotide sequences of all *opa* genes using MAFFT v7.520 [35] with a gap opening penalty of 4 and a gap extension penalty of 1. The nucleotide sequence of FA1090 *opa1* was compared to the sequences in Bhat et al. [19] to identify the semivariable, hypervariable 1, and hypervariable 2 region sequences. The variable regions in the other *opa* sequences were determined based on where they aligned to FA1090 *opa1*.

#### Calculation of k-mer distance

The k-mer distances between sequences in each of the semivariable, hypervariable 1, and hypervariable 2 regions were calculated using MASH v2.3 [46] with a k-mer size of 6 for the semivariable region and 7 for the hypervariable 1 and hypervariable 2 regions. These k-mer sizes were chosen so that the probability of observing a random sequence of length k in a random sequence the length of each region was 0.01. The p-value is the probability of observing a given k-mer distance (or less) under the null hypothesis that both sequences are random collections of k-mers.

#### Clustering

The variable region sequences were clustered using the Markov Cluster Algorithm (MCL) v14.137 [48]. We constructed a distance matrix using the -log10 transformed p-value from MASH and values larger than 200 were set to 200. We made these transformations following the approach in TRIBE-MCL, which clusters proteins using -log10 transformed e-values from BLAST [49]. The distance matrix was input into MCL, which then performs multiple rounds of expansion and contraction to amplify large values and reduce small values of the matrix. The number of rounds of expansion and contraction is set by the inflation parameter. We ran the clustering using a range of inflation parameters: 1.4, 2, 4, 6, 8, 10, 12, and 14. We chose the inflation parameter that gave a stable clustering, inspired by TRIBE-MCL. A stable clustering was defined as having a percentage difference in clustering with the next lowest inflation parameter as less than 1% for the semivariable and hypervariable 2 regions. For the hypervariable 1 region, we relaxed the definition for a stable clustering as having less than a 5% difference in clustering with the next lowest inflation parameter due to the large number of clusters. The percentage difference in clustering is calculated as the sum of the number of nodes (sequences) that must be exchanged to transform the two clusterings into each other (also called the split/join distance) divided by two times the number of sequences. The motivation for this approach is that TRIBE-MCL uses convergence of outputs to determine the inflation parameter.

The cluster sequence logos were visualized by aligning the nucleotide sequences in each cluster using MAFFT v7.520 [35] with the default parameters and plotting the sequence logos with Logomaker v0.8 [50].

### Ancestral state reconstruction

#### Dated phylogeny

Pseudogenomes were generated as described above but with the reference genome being one randomly selected genome from the subtree. Regions of the genome under recombination were detected using Gubbins v3.3.4 [44] (GTR substitution model, RAxML Next Gen tree builder, 20 iterations) and masked. The recombination-masked alignment was used to build a dated phylogeny in BEAST2 v2.7.5 [51], for all isolates that had date of collection information. When the date was given as a year, we set the date to January 1 of that year. We used a GTR substitution model with gamma rate heterogeneity (4 categories), a strict clock model, a coalescent constant population, and 100,000,000 chain length. Chains were inspected to confirm good mixing (effective sample size greater than 200).

#### Ancestral state reconstruction

We performed ancestral state reconstruction for each of the loci and variable region cluster types separately on the dated phylogeny using PastML v1.9.49 [52] with the default parameters. Isolates that did not have an intact *opa* at a locus were not included for the inference at that locus. If there were no changes to the variable region cluster type at a locus then the average rate of cluster type changes was set to 0.

### Phylogenies

Mutation-scaled phylogenies were visualized in iTOL v6.9.1 [53] using midpoint rooting. The time-scaled phylogeny was visualized in FigTree v1.4.4 (http://tree.bio.ed.ac.uk/software/figtree/).

### Statistical analyses

Statistical analyses and plotting were done in Python v3.13.1.

## Data availability

The Nanopore sequencing reads and assembled complete genomes generated in this study are available at the European Nucleotide Archive under study accession PRJEB106623: http://www.ebi.ac.uk/ena/browser/view/PRJEB106623.

## Code availability

Our code for performing the analyses has been compiled into a Snakemake [54] pipeline for reproducibility. The code is available in GitHub (https://github.com/qinqin-yu/gonococcus_opa_diversity).

## Results

### A diverse set of 219 complete genomes of *N. gonorrhoeae*

Because long-read only assemblies using older sequencing technologies have a higher error rate, we filtered the publicly available complete genome assemblies to include only those assembled using long-read and short-read data, which yielded 132 complete genomes (**Table S1**). To complement the publicly available complete genomes of *N. gonorrhoeae*, we sequenced an additional 87 isolates from across the *N. gonorrhoeae* phylogeny using Oxford Nanopore long-read sequencing (**Figure 1, Table S2**). In total, this gave 219 complete genomes.

**Figure 1:**
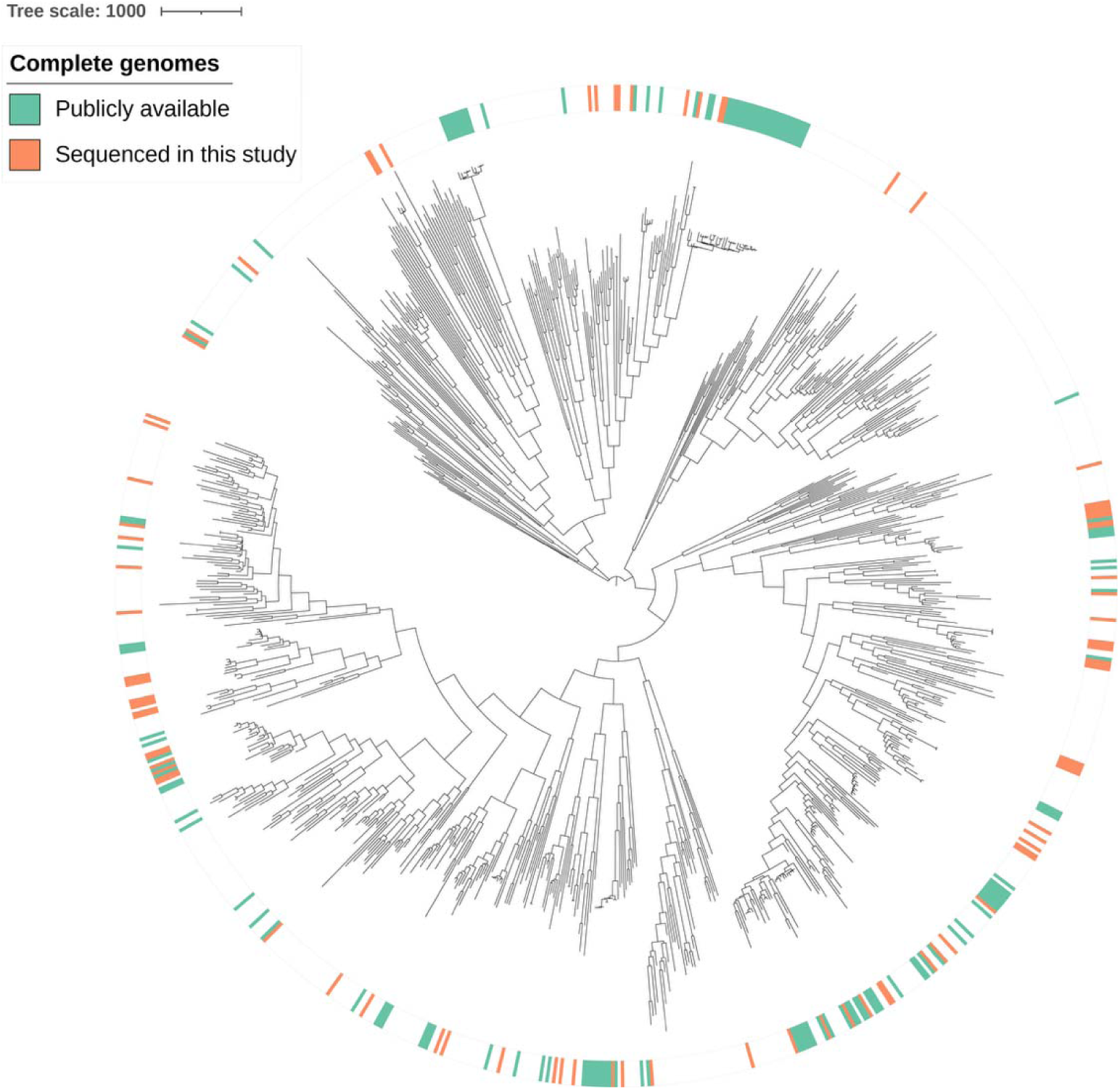
A diverse set of *N. gonorrhoeae* complete genomes. Recombination-masked phylogeny of the publicly available complete genomes (generated with long- and short-read sequencing data, green) and the new complete genomes generated in this study (generated with Oxford Nanopore sequencing, orange), in the context of a representative global collection of *N. gonorrhoeae* isolates (generated with short-read sequencing data). The tree scale indicates the number of recombination-masked SNPs.

Comparison of three long-read only assembly methods (see Supplementary Methods) on two isolates whose genomes differed by 6648 SNPs showed that Autocycler performed the best, with no changes to the *opa* sequences for read depths of 21-140x (**Figure S*1*** and Supplementary Information). Polishing the Autocycler assemblies with short-read data did not change the *opa* sequences at any read depth, showing that the long-read data was sufficient to reconstruct the *opa* sequences. Further testing on a larger set of 11 diverse isolates indicated that varying read depths resulted in differences in *opa* sequence for only 4 out of 119 *opa* sequences in long-read only assemblies with Autocycler (3 SNPs and one *opa* that was undetected or had large numbers of sequence differences; see **Figure S*2*** and Supplementary Information).

### Identification of *opa* genes

Across the 219 complete genomes, we identified 2,359 full length *opa* genes. To determine if we missed any *opa* sequences, we compared our approach to three standard strategies to identify homologous genes: BLAST, Prokka, and pangenome clustering of the Prokka annotations. Using BLAST to search for the conserved region of the *opa* gene did not identify any additional intact *opa* genes compared to our method, but it did identify 35 additional truncated *opa* genes and 2 *opa* genes with a mutation in the start codon (ACG instead of ATG). Our method identified 8 *opa* genes not found by BLAST that were divergent from other *opa* sequences. Gene annotations using Prokka identified the same 37 truncated *opa* genes or *opa* genes with a mutation in the start codon found by the BLAST search, but no additional genes. Our method identified 2 additional *opa* genes that were not annotated by Prokka. Pangenome clustering identified the 37 truncated *opa* genes or *opa* genes with a mutation in the start codon found by the BLAST search and one additional truncated *opa*. Our method identified 7 *opa* genes not found in the pangenome clusters, including the 2 *opa* genes that were not annotated by Prokka. In summary, our method of searching for *opa* genes identified the most comprehensive set of *opa* genes from the genomes and, to our knowledge, did not miss any intact *opa* genes.

### Consistent local genomic positioning of *opa* reveals patterns of copy number variation

We next determined the number and locations of *opa* genes in each genome. Out of 219 genomes, 5 genomes (2.3%) had 9 *opa* genes, 41 genomes (18.7%) had 10 *opa* genes, 172 genomes (78.5%) had 11 *opa* genes, and 1 genome (0.5%) had 12 *opa* genes. The genome with 12 *opa* genes (WHO_T_2024) contained a duplication of 47,339 nucleotides with 2 SNPs that led to the exact duplication of one of the *opa* genes. Based on the genomes in this study, we observed *opa* losses a handful of times across the phylogeny (Figure 2a). In addition, there were 35 sequences that contained part of an *opa* gene. Of these, 28 occurred in a phylogenetically related cluster of isolates and were missing the last 260 nucleotides of the gene, including the hypervariable 2 region, in the context of an approximately 400 nucleotide deletion that also included 140 nucleotides of sequence downstream of the *opa* gene. These 28 clustered, truncated *opa* sequences were similar but not identical and occurred in similar locations in the genome. There were also 2 sequences with a mutation in the start codon (ACG instead of ATG) that otherwise were full length *opa* genes. All but one of the truncated *opa* sequences and both *opa* sequences with a mutation in the start codon occurred in genomes that had fewer than 11 intact and full length *opa* genes. If all genomes were to have at least 11 *opa*, then there would be at least 51 missing full length *opa* across the dataset, 37 of which are accounted for here.

**Figure 2:**
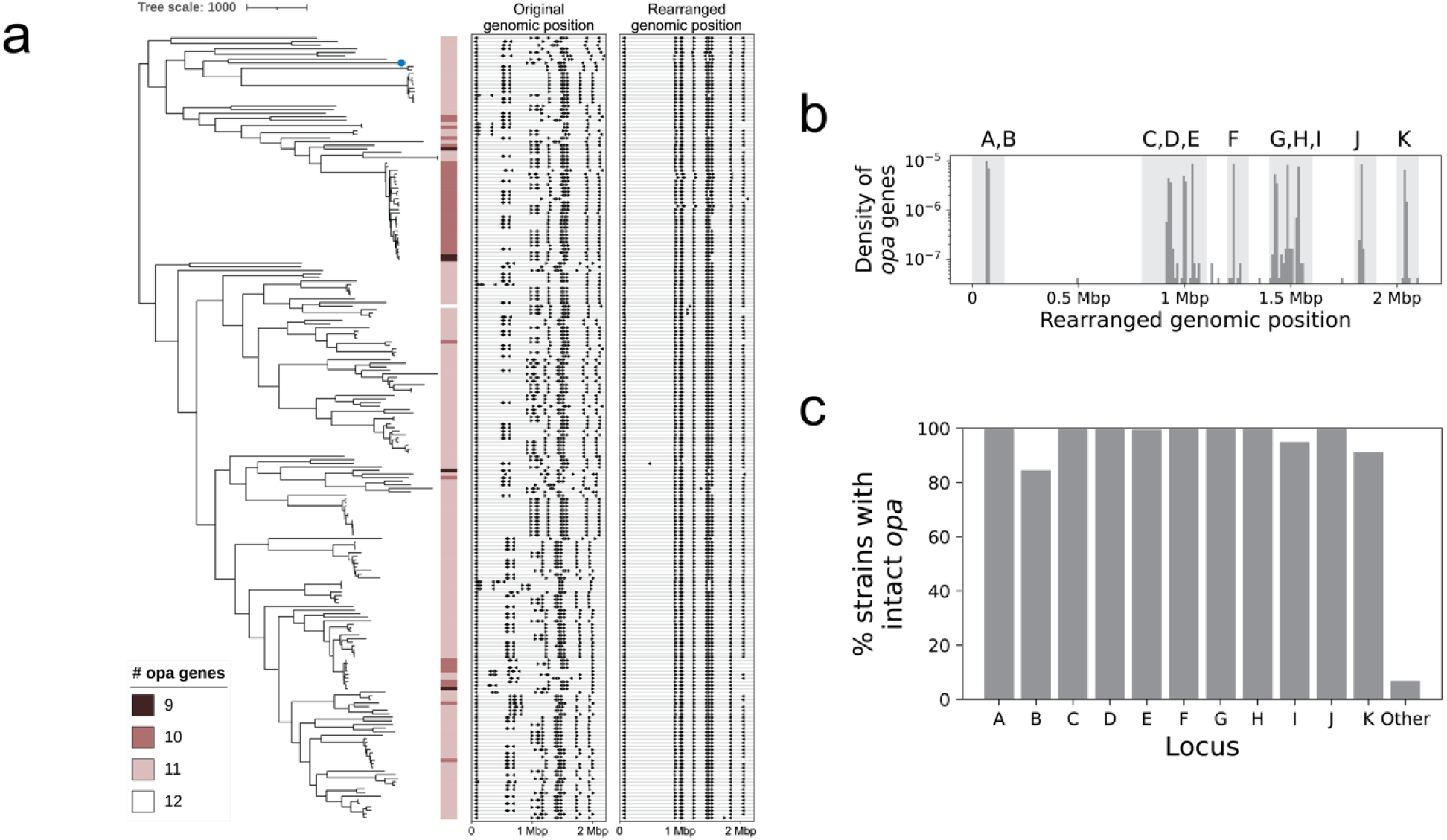
Consistent local genomic positioning of *opa* reveals patterns of copy number variation. (a) Recombination-masked phylogeny of complete genomes annotated with the number of intact *opa* genes (contain start codon, stop codon, and N-terminus) in each genome. The genomic locations of the *opa* genes are shown in the middle panel and the rearranged genomic locations of the *opa* genes are shown in the right panel. The reference genome (FA1090) is shown with a blue dot on the phylogeny. The tree-scale indicates the number of recombination-corrected SNPs. (b) The distribution of number of *opa* genes by rearranged genomic position. The loci were named *opaA* to *opaK* based on the FA1090 genomic order. *opa* genes that fell within the light-gray shaded background regions were assigned the corresponding name(s). Where multiple *opa* genes occurred in proximity (for example, A and B; C, D, and E; or G, H, and I), the *opa* genes within each genome were assigned in the order that they appeared in the light-gray shaded region (i.e., the first *opa* gene is C, the second *opa* gene is D, and the third *opa* gene is E). This was done to account for slight shifts in *opa* location across genomes. (c) *opa* loci occupancy.

The genomic locations of *opa* genes were variable across genomes (**Figure S*3***), with a mean distance of 59,460 nucleotides between *opa* gene loci across strains. We observed that the pattern of genomic organization of *opa* loci clustered phylogenetically (Figure 2a). To test whether differences in *opa* locations were due to large-scale genomic rearrangements, we used alignments to the reference strain FA1090 to detect and then correct for such rearrangements (**Figure S*4***). We found 2,331/2,359 (98.8%) *opa* genes within locally collinear alignment blocks with the FA1090 genome. The mean distance between *opa* locations in the rearrangement-corrected assemblies decreased to 2,886 nucleotides (Figure 2a**, Figure S*3***). The consistency of *opa* locations in the rearrangement-corrected assemblies allowed us to define 11 loci, which we named in the order that they appeared in FA1090 from *opaA* to *opaK* (Figure 2b). Loss of full-length *opa* genes occurred most often in *opaB* (16% of all isolates), *opaK* (9% of all isolates), and *opaI* (5% of all isolates) (Figure 2c). There were 15/2,359 (0.6%) *opa* that did not fall within these 11 loci.

Of the 2,359 *opa* genes, 156 (6.6%) had a frameshift mutation after the end of the coding repeat sequence, leading to a stable premature stop codon independent of phase variation. This included at least one *opa* gene in 51/132 (39%) of publicly available genomes and 29/87 (33%) genomes sequenced in this study. None of the superseded WHO reference strains [55] had these frameshift mutations. For the genomes sequenced in this study, there was a negative correlation between the sequencing coverage and the number of *opa* with these frameshift mutations (**Figure S*5*a**, Spearman correlation coefficient −0.45, p=10^-5^), suggesting that some of these frameshift mutations may be due to sequencing error. The premature stop codons were concentrated in three locations in the *opa* gene (**Figure S*5*b**) and closer inspection of the nucleotide sequences revealed that variations in the semivariable region sequence upstream of the premature stop codon led to the frameshift mutations.

### Diversity of *opa* alleles within genomes, across genomes, and compared with genome diversity

*opa* genes had almost as much within-isolate diversity as between-isolate diversity, as measured by pairwise amino acid sequence identity (Figure 3a). The number of distinct *opa* allele types per isolate, as defined by having less than 95% amino acid identity, ranged from 3 to 10, with a mean of 7 (Figure 3c). The Shannon diversity of *opa* types in each isolate ranged from 1.03 to 2.27 with a mean of 1.84 (**Figure S*6***). There were more instances of similar *opa* genes in the same isolate than in different isolates, and 96.8% (212/219) of isolates had at least one pair of *opa* genes with more than 95% amino acid sequence identity (Figure 3b). One isolate (NG250) had 7 *opa* genes with >95% amino acid sequence identity and three isolates (CT213, TUM15748, and WHO_T_2024) each had 6 *opa* genes with >95% amino acid sequence identity. Thirteen isolates had 4 pairs of *opa* genes with >95% amino acid sequence identity (**Figure S 7**). *opa* sequence similarity correlated with genome similarity (Spearman correlation coefficient 0.24, p < 0.001) (Figure 3c). In pairs of non-identical isolates, the number of sites in the *opa* genes that differed was 13-3,294x higher (mean: 74x higher) than the number of whole genome sites that differed, showing that *opa* genes are substantially more diverse than the rest of the genome (Figure 3d).

**Figure 3:**
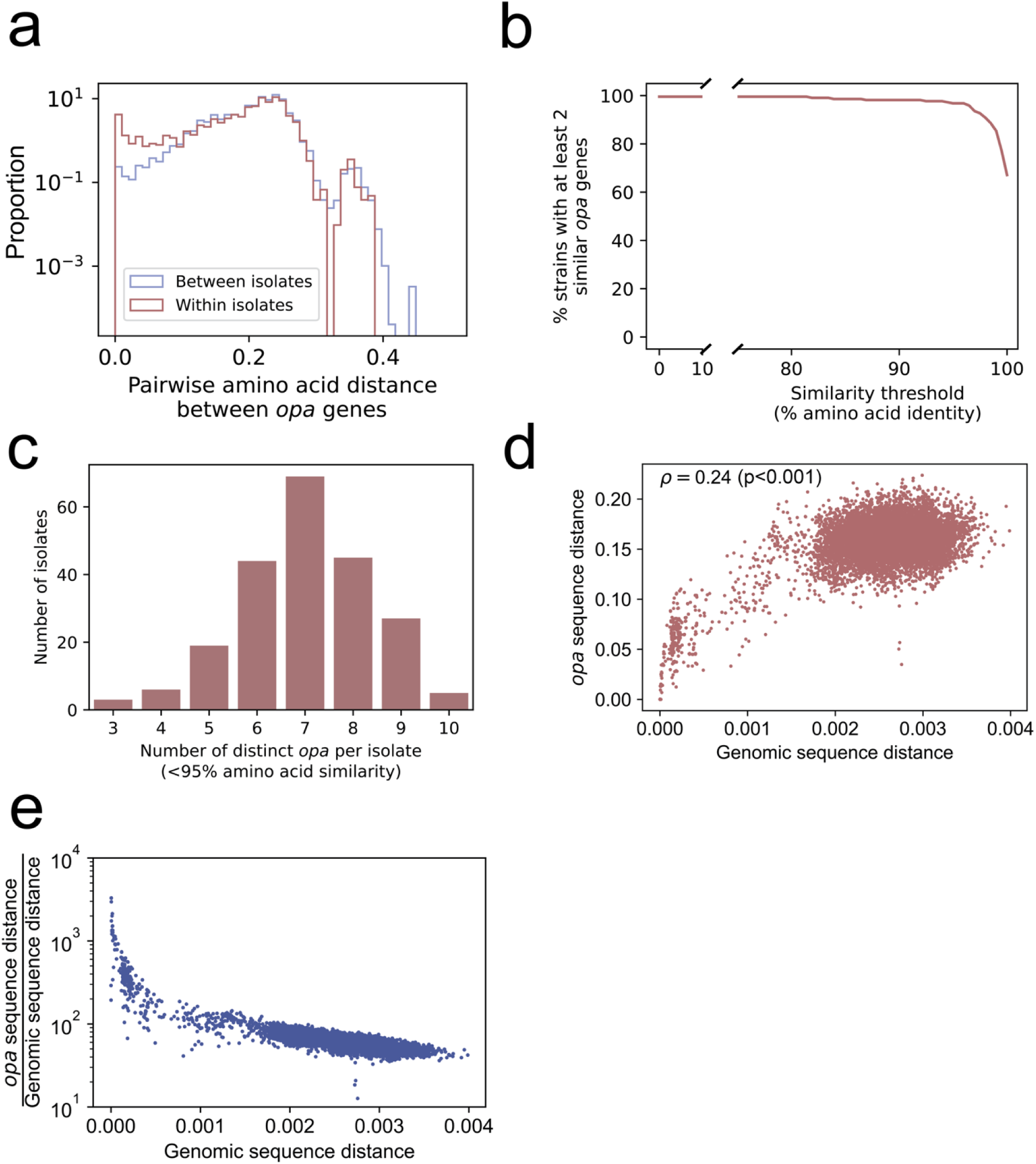
Diversity of *opa* alleles within genomes, across genomes, and compared with genome diversity. (a) The distribution of the pairwise amino acid distance between *opa* genes in the same isolate (red) and in different isolates (blue). The distribution of within-isolate *opa* diversity was similar to the distribution of between-isolate *opa* diversity, except for the presence of more near-identical *opa* genes in the same isolate. (b) The percentage of isolates with at least 2 *opa* sequences above a given amino acid sequence identity. Nearly all (97.2%) isolates have at least 2 *opa* sequences with >95% amino acid sequence identity. (c) The distribution of the number of distinct *opa* allele types per isolate, defined as less than 95% amino acid sequence identity. (d) Comparison of *opa* sequence distance and genomic sequence distance in pairs of complete genomes with the same number of *opa* genes. Each point represents one pair of complete genomes with the same number of *opa* genes. A distance of 0 indicates identical, and a distance of 1 indicates every site is different. The Spearman correlation coefficient and p-value are shown in the upper left. (e) The same data as in (d) but with the *opa* sequence distance divided by the genomic sequence distance on the y axis.

### Fewer *opa* genes are in frame than expected by chance

193/219 (88%) isolates had at least 1 *opa* gene in frame (Figure 4). The mean number of *opa* genes in frame across all isolates was 1.7 and the maximum of *opa* genes in frame in any isolate was 5. The mean number of in-frame *opa* genes per isolate was significantly lower (p<0.001, Wilcoxon signed-rank test) than expected from a simple null model where each *opa* has a 1/3 chance of being in frame (mean of 3.4 in-frame *opa* genes per isolate in null distribution). *opa* genes that were more than 95% identical in amino acid sequence to at least one other *opa* in the same isolate were 1.54 times less likely to be in frame than other *opa* genes in the isolate (p<10^-4^) (**Figure S 8**).

**Figure 4:**
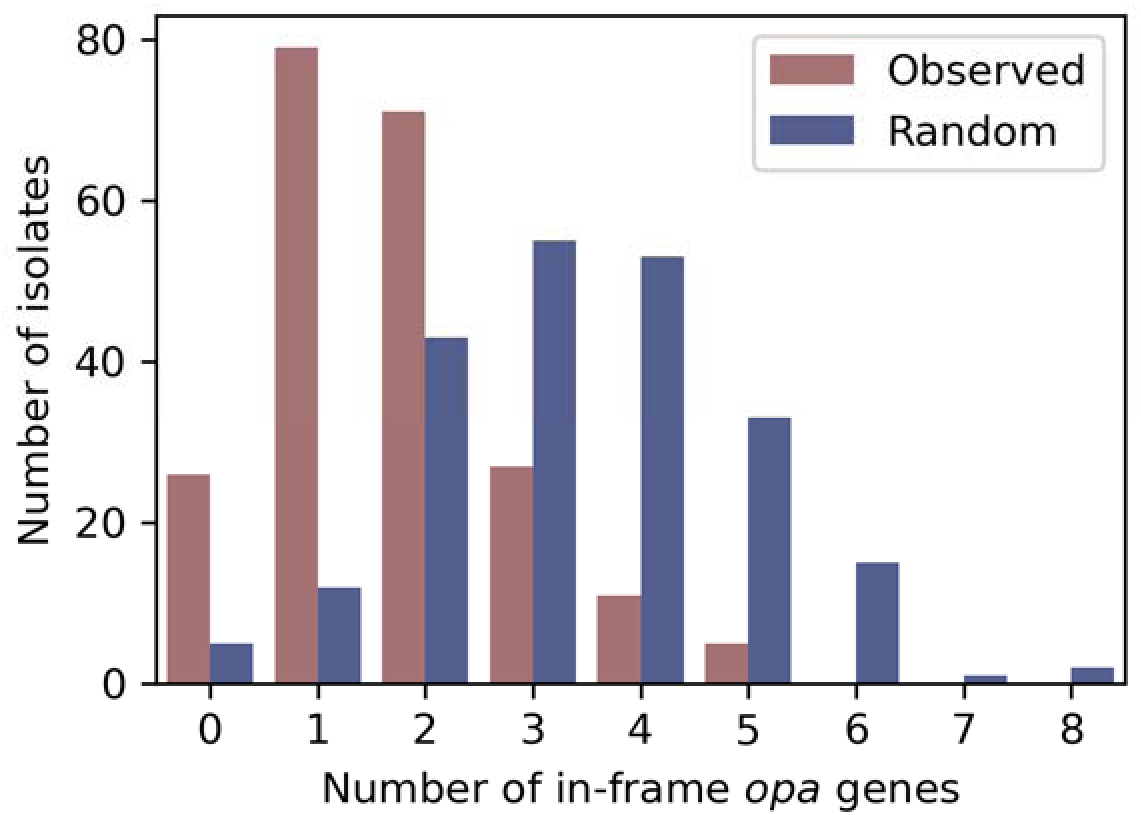
Fewer *opa* genes are in frame than expected by chance. The distribution of in-frame *opa* genes per isolate.

### N. gonorrhoeae *opa* are phylogenetically distinct from *Neisseria* species *opa*

As *N. gonorrhoeae* can gain new genetic diversity through interspecies mosaicism [56,57], we evaluated the possible contribution of other *Neisseria* to *N. gonorrhoeae opa* diversity (**Figure S 9**). Our approach to searching for the *opa* genes in *Neisseria* species using conserved sequences identified the same number of intact *opa* genes as BLAST, and BLAST identified one additional truncated *opa* gene. Most *N. meningitidis* isolates had 4 full length *opa* (115/136 isolates), and more rarely 3 full length *opa* (19/136 isolates) or 5 full length *opa* (2/136 isolates).

*N. flavescens* and *N. lactamica* isolates had 2 full length *opa* (1 isolate of *N. flavescens* and 4 isolates of *N. lactamica*), and *N. polysaccharea* had 1 full length *opa* (1 isolate) (Figure 5a). Multiple *Neisseria* species had partial *opa* genes that contained the C-terminal region of the *opa* gene only: *N. animalis* (2 isolates with 2 partial *opa* each), *N. arctica* (1 isolate with 2 partial *opa*)*, N. brasiliensis* (1 isolate with 1 partial *opa*)*, N. sicca* (1 isolate with 1 partial *opa*, 2 isolates with no *opa*)*, and N. zalophi* (1 isolate with 1 partial *opa*). Additionally, 11/136 *N. meningitidis* isolates had a partial *opa* which all have the same sequence (including the upstream region)*. N. gonorrhoeae opa* were phylogenetically distinct from *opa* from other *Neisseria* species, except for one allele, which was found in 8 *N. gonorrhoeae* isolates. This allele clustered with, and its sequence was similar to, commensal *Neisseria opa* (Figure 5b).

**Figure 5:**
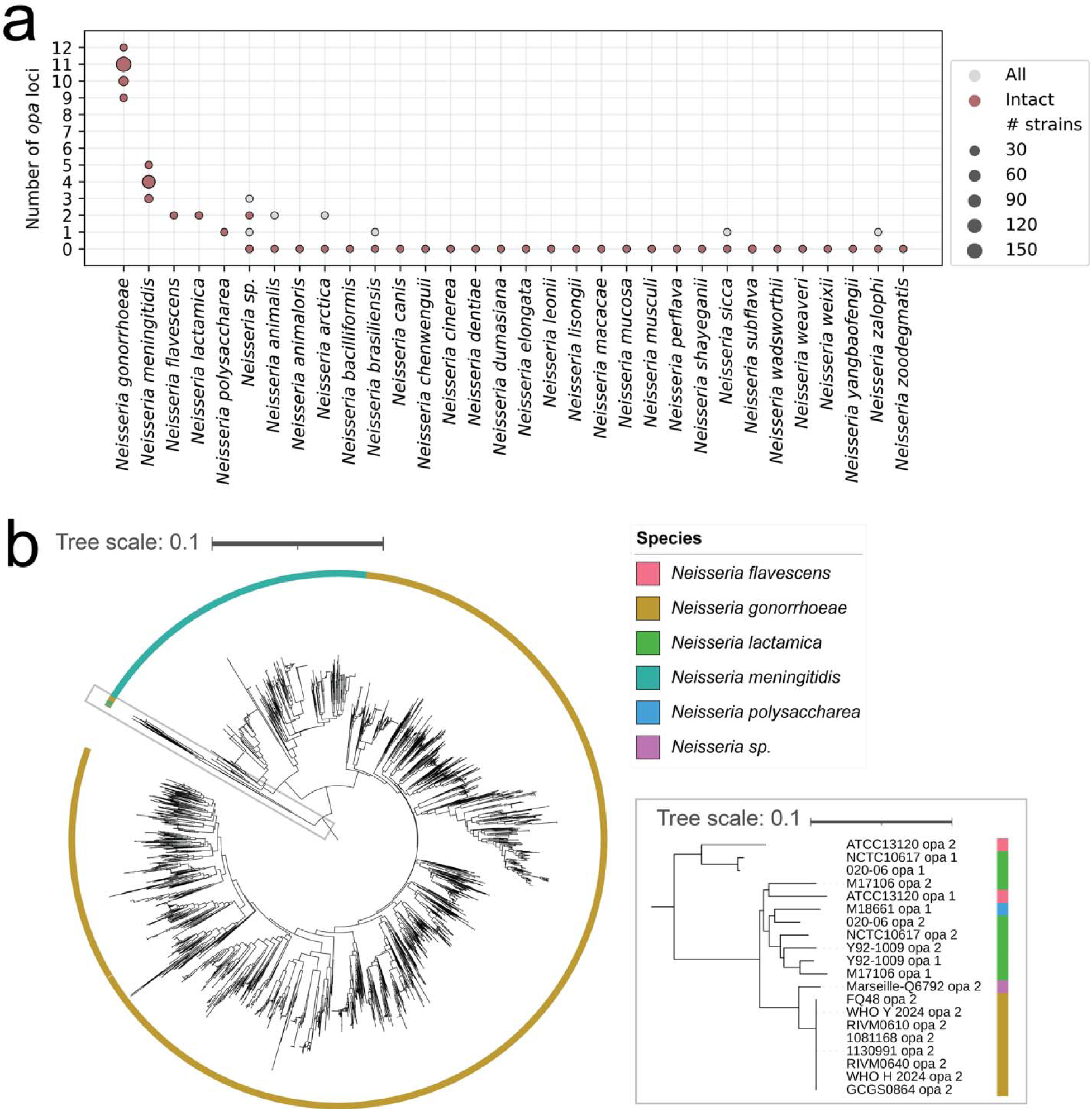
*N. gonorrhoeae opa* are phylogenetically distinct from other *Neisseria* species *opa*. (a) The distribution of the total (partial and intact) and intact number of *opa* loci by species. (b) k-mer distance-based neighbor-joining phylogeny of the *opa* sequences in all *Neisseria* species. *N. gonorrhoeae opa* are phylogenetically distinct from *opa* sequences from other *Neisseria* species, with the exception of one allele that clustered with other *Neisseria* species *opa* (inset, which shows zoom of the boxed section of the phylogeny). The tree scales indicate the k-mer distance.

### The rate of *opa* evolution is dependent on the genetic locus

To estimate the rate of evolution, we first clustered *opa* by sequence similarity (**Figure S*10***). Because of known recombination-mediated shuffling [10,11] of the semivariable, hypervariable 1, and hypervariable 2 sequences (**Figure S*10*a**), we clustered each of these regions separately using k-mer distances and a network clustering approach (**Figure S*10*b**). The clustering resulted in 6 semivariable region clusters, 21 hypervariable 1 region clusters, and 9 hypervariable 2 region clusters (**Figure S*10*c**). The nucleotide k-mer distances for sequences in the same cluster are lower than the distances for sequences in different clusters (**Figure S*11***), and examination of the sequences shows that each cluster has a distinct motif (**Figure S*12*-14**).

We reconstructed the ancestral states of the *opa* cluster types on a dated phylogeny to estimate the rate of *opa* evolution. We focused on the only clade in the phylogeny of complete genomes that had densely sampled isolates (branch length less than 200 recombination-corrected mutations and more than 20 samples) (Figure 6a). The inferred substitution rate, 3.1×10^-6^ [1.7×10^-6^, 4.4×10^-6^] substitutions per chromosome per year, fell within the rate expected for *N. gonorrhoeae* [58]. The rate of cluster type changes was substantially higher for *opaK* compared to the other *opa* loci (Figure 6b). The cluster types that appeared in *opaA* and *opaK* were similar, showing that the difference in the rates of cluster type changes between the two loci was due to different rates of shuffling between the same cluster types (**Figure S*15***). *opaK* was not significantly more in frame than the other *opa* loci (**Figure S*16***).

**Figure 6:**
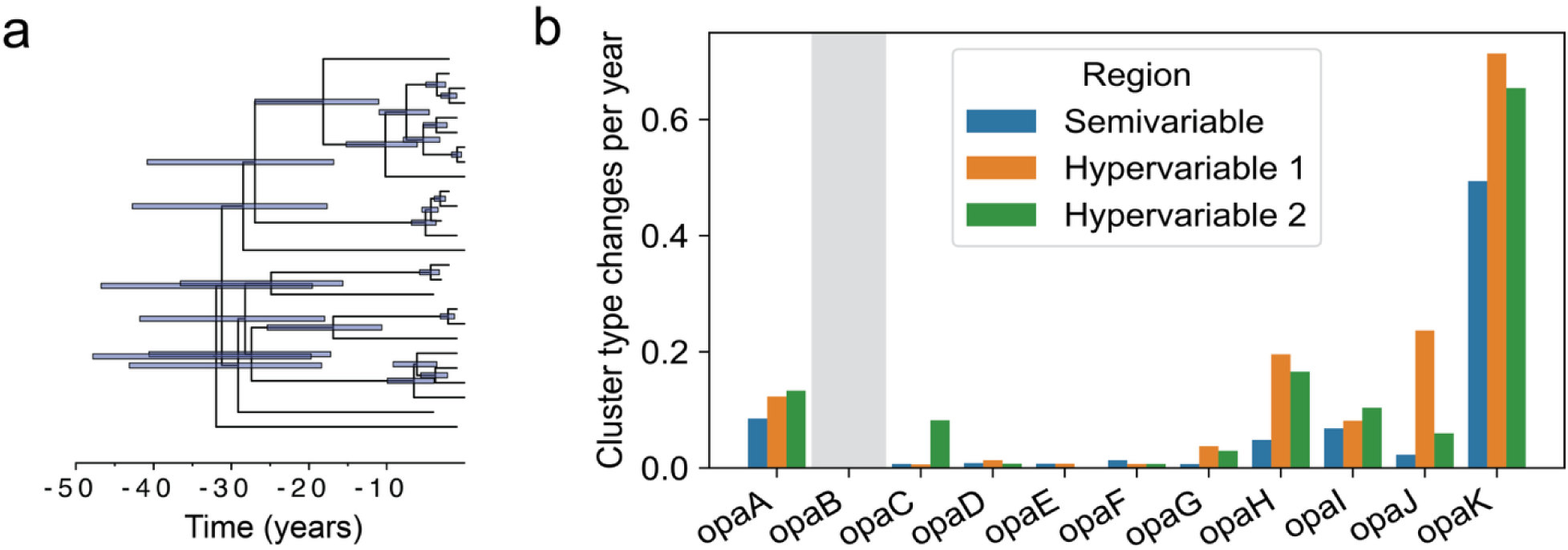
The rate of *opa* evolution is dependent on the genetic locus. (a) Dated phylogeny of the clade in Figure 2a with densely sampled isolates (branch length less than 200 recombination-corrected mutations and more than 20 samples). Error bars represent the 95% highest posterior density of the node heights. (b) Rate of semivariable, hypervariable 1, and hypervariable 2 cluster type changes by *opa* locus for subtree depicted in (a). The gray shaded region denotes that *opaB* was missing from all isolates in this subtree. In other loci, there were no more than 2 isolates missing each *opa*.

### Correlation of hypervariable 1 and hypervariable 2 region alleles

Using the cluster typing, we were also able to assess whether certain hypervariable 1 and hypervariable 2 alleles appeared together more often than expected by chance (**Figure S*17***).

We accounted for isolate sampling and population structure by randomly sampling 1 isolate per whole genome cluster 100 times. For each subset of representative isolates, out of 100 randomizations of the data, the actual data always showed higher levels of association between hypervariable types than the randomized data.

## Discussion

### Quantitative assessment of *opa* diversity, expression, and evolution

Prior to this study, there were several open questions about *opa* genes in *N. gonorrhoeae*: how are *opa* genes organized in the genome? Are there patterns in their diversity and expression? How much *opa* diversity is there in the population? How do *opa* evolve? Here, we have advanced our understanding for each of these questions using a new diverse dataset of complete *N. gonorrhoeae* genomes.

### Genomic rearrangements shuffle opa position

Genomic rearrangements shuffle *opa* position in the genome, but by accounting for them, we were able to assign consistent *opa* loci. Genomic rearrangements in closely related isolates happened very rarely, but there is space for the locus assignment approach to be updated if we see more examples of genomic rearrangements in closely related isolates.

### Gain and loss of opa occurred repeatedly

While most isolates had 11 *opa* genes, we found a handful of instances of *opa* loss and gain. The loss events occurred across the phylogeny. In some instances, we saw examples of the gene loss events passed on to progeny. More than 50% of these loss events were explained by truncated *opa* genes. We were unable to identify the mechanism for the other loss events.

Furthermore, in intact *opa* sequences, we observed frameshift mutations occurring downstream of the coding repeat sequence leading to a premature stop codon, but we were unable to determine how many of these frameshift mutations were due to sequencing error. In one isolate, we identified 12 *opa* genes, which seemed to be due to a recent genomic duplication event that exactly duplicated another *opa* gene in the genome. The fact that we only observed one isolate with 12 *opa* suggests that the expansion of *opa* number is rare.

### Within-isolate opa diversity is generally high, with some exceptions

The diversity of *opa* alleles within isolates was generally high, with the average isolate having 7 distinct *opa* alleles, but there were some isolates with as few as 3 distinct *opa* alleles. This suggests that there is some pressure to have diverse *opa* alleles within an isolate. At the same time, almost all isolates had at least two near-identical *opa* alleles, possibly as a result of gene conversion, suggesting there may be a balance between maintaining diversity and maintaining function while diversifying.

### Fewer opa expressed than expected by chance

Across the population, fewer *opa* appeared to be expressed based on the sequence than expected by chance. Furthermore, near-identical *opa* genes in the same isolate tended to be out of frame more often than unique *opa* genes. Our results suggest mechanisms other than chance alone in guiding the patterns of phase variation, for example, regulation of changes in repeat number or selection against isolates with many *opa* alleles being expressed. While these mechanisms are yet unknown, previous work suggested that promoter strength may play a role in regulating *opa* phase [59] and there may be selection against *opa* expression in certain environments, for example the presence of progesterone [60–62].

### opa diversity is correlated with and higher than genome diversity

*opa* genes are on average 74 times more diverse than the rest of the genome. More closely-related isolates have more similar *opa* sequence repertoires. In the context of reinfections, our results suggest that when reinfections occur with the same or a closely related isolate, there will be a similar *opa* repertoire. We speculate that if there is immunological memory to the *opa* that are expressed, then different *opa* are turned on in subsequent reinfections, an idea that is also supported by evidence of variation in *opa* expression over the course of single infections in human challenge studies [63,64]. When reinfections occur with different isolates, there will likely be different *opa* alleles expressed.

### Interspecies recombination does not appear to play a large role in generation of opa diversity

*N. gonorrhoeae opa* allelic diversity did not overlap with *opa* from other *Neisseria* species. While sequencing more complete genomes may lead to more observations of interspecies mosaicism, the separate clustering of *N. gonorrhoeae* and *N. meningitidis opa* in our dataset and in Malorny et al. [9] suggests phylogenetic separation of *opa* sequences between species. The separate clustering of *Neisseria* species *opa* may be due to the different functional constraints of host receptor binding or differences in flanking regions of *opa* genes from different species that prevent homologous recombination from occurring between species.

### Variation in rate of opa variation across loci

*opaK* varies more rapidly than other loci, but it is not significantly more in frame in our dataset. The differences in rates of evolution by *opa* locus may be due to variability in the accessibility of each locus for recombination or other diversity generating processes (i.e., structural accessibility). The changes in *opaK* tend to reflect recombination among existing *opa* types, suggesting that diversity is mostly generated through shuffling of existing sequences. Bilek et al. [21] suggested a possible connection between *opa11* (*opaK* in this study) variation and the neighboring *pilE* Pilin expression gene variation, but noted this may not be causal.

### Hypervariable region 1 and 2 alleles are more correlated than expected by chance

Hypervariable region 1 and 2 alleles in *N. meningitidis* are more correlated than expected due to chance [65]. We showed that this phenomenon also occurs in *N. gonorrhoeae*. The non-random association of hypervariable alleles is consistent with predictions of a mathematical model of strong immune selection [66], physical linkage on the chromosome, and selection for particular combinations of hypervariable alleles for functional reasons (i.e., to bind to specific host receptors)[67].

### Limitations of our study

There are several limitations to our study. First, the number of complete genomes sequenced was limited, possibly missing some *opa* diversity. Second, gonococcal passaging during culturing may have changed the phase of *opa* genes. Culture-independent sequencing that resolves *opa* phase *in vivo* will add to our understanding of patterns of *opa* expression. Third, there is limited data on *opa* function, with only a few *opa* sequences characterized for their binding to host receptors and antibody recognition [15]. While the clustering nomenclature for *opa* that we developed is entirely sequence based, we expect closely related sequences to have more closely related functions on average. In the future, functional data can be incorporated to refine the *opa* clustering definitions. Fourth, we were limited in the estimation of the rate of evolution to only a single densely sampled subtree. There may be differences in the rate of *opa* evolution across genomic backgrounds.

### Model

Taken together, our results support a model of *opa* variation where on short timescales phase variation modulates the translation of highly diverse *opa* alleles, typically with very few *opa* being expressed at one time. On long timescales, recombination, gene conversion, mutation, and gene gain and loss lead to changes in the *opa* repertoire, with the rate of these processes dependent on the genetic locus (Figure 7). The rapid rate of phase variation and diversification will likely lead hosts to be exposed to different Opa types upon reinfection and may allow antigenic escape in vivo in chronic or undiagnosed infections.

**Figure 7:**
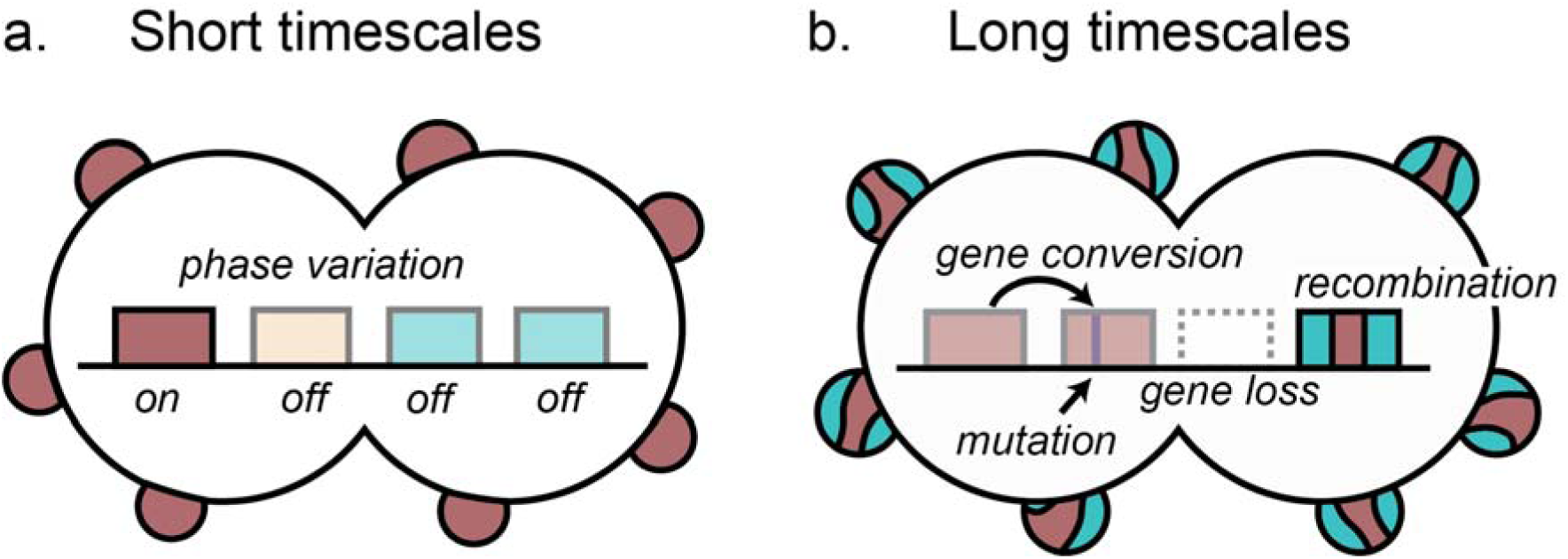
Model of *opa* variation and evolution supported by our results. (a) On short timescales, phase variation modulates the expression of highly diverse *opa* alleles within isolates, typically with very few *opa* being expressed at one time. (b) On long timescales, recombination, gene conversion, mutation, and gene gain and loss within species in a locus-dependent manner lead to changes in the *opa* repertoire.

## Supporting information

Table S1 references

Table S2 references

Table S3 references

## Acknowledgements

We are grateful to Eric Neubauer Vickers for helpful discussions and comments on the manuscript. We are grateful to the Grad lab and the Center for Communicable Disease Dynamics for helpful discussions. This project received support from the American Sexually Transmitted Diseases Association (to YHG and QY), NIH R21 AI172369 (to YHG), and NIH T32 AI007535 (to QY). The computations in this paper were run on the FASRC Cannon cluster supported by the FAS Division of Science Research Computing Group at Harvard University.

## Supplementary figures

**Figure S1:**
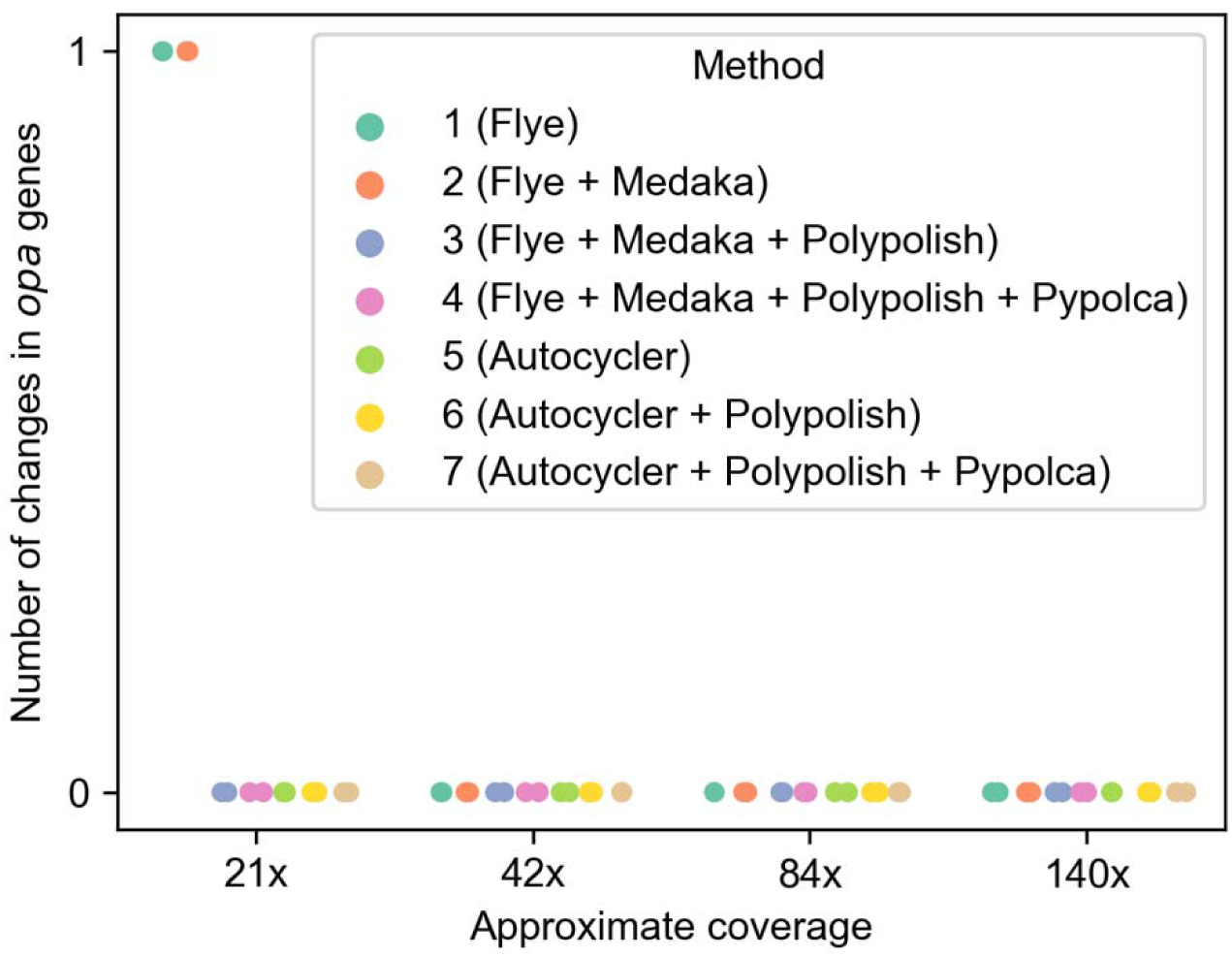
Comparison of *opa* sequences across long-read assembly and polishing methods. The number of changes in *opa* genes at 4 read coverage levels using 7 different assembly and polishing procedures. There are two points of the same color in each coverage level, indicating two different isolates. The changes at 21x for methods 1 and 2 include one SNP in one isolate’s genome and one single base insertion in the other isolate’s genome.

**Figure S2:**
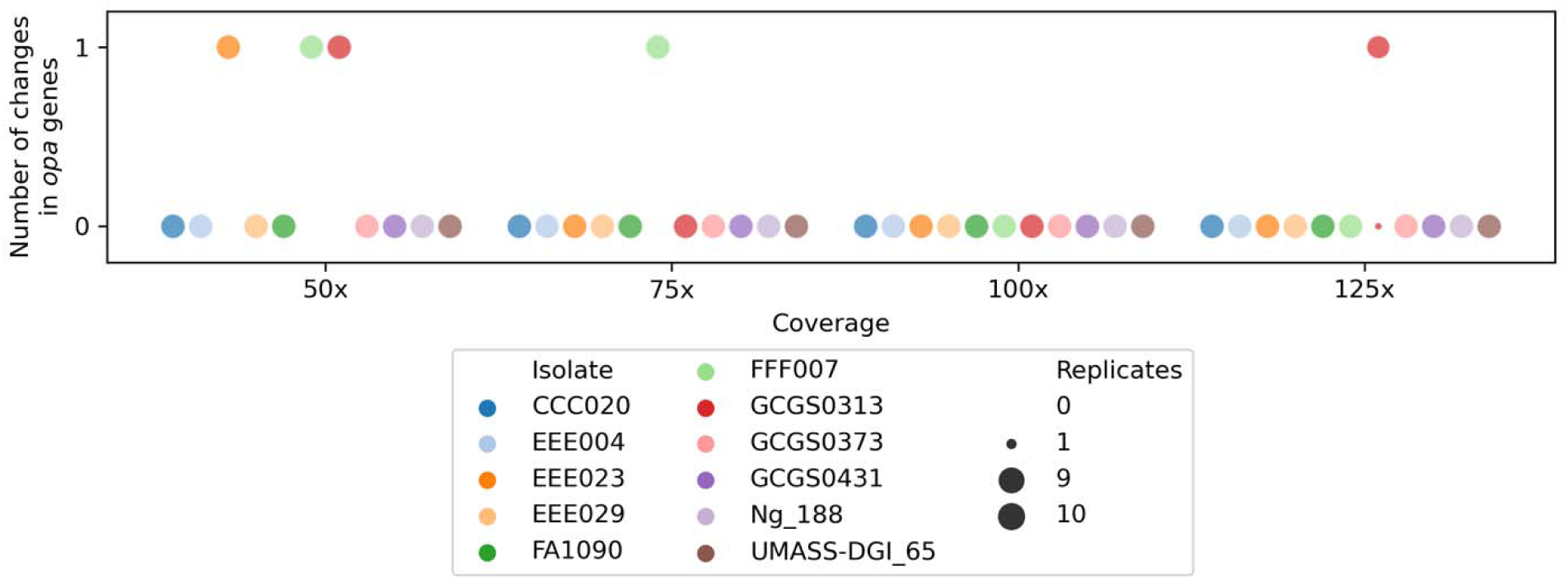
More extensive comparison of *opa* sequences in Autocycler assemblies across 11 diverse isolates at 4 read coverage levels. The number of changes in *opa* genes at 4 read coverage levels using Autocycler. For each isolate, the reads were randomly subsampled 10 times at each read coverage (replicates) and an assembly was created with the subsampled reads using Autocycler. The size of the point indicates the number of replicates. The changes at 50x coverage include one SNP in two separate genomes and one undetected *opa*, the change at 75x was multiple sequence differences in one genome, and the change at 125x coverage was one SNP in one genome.

**Figure S3:**
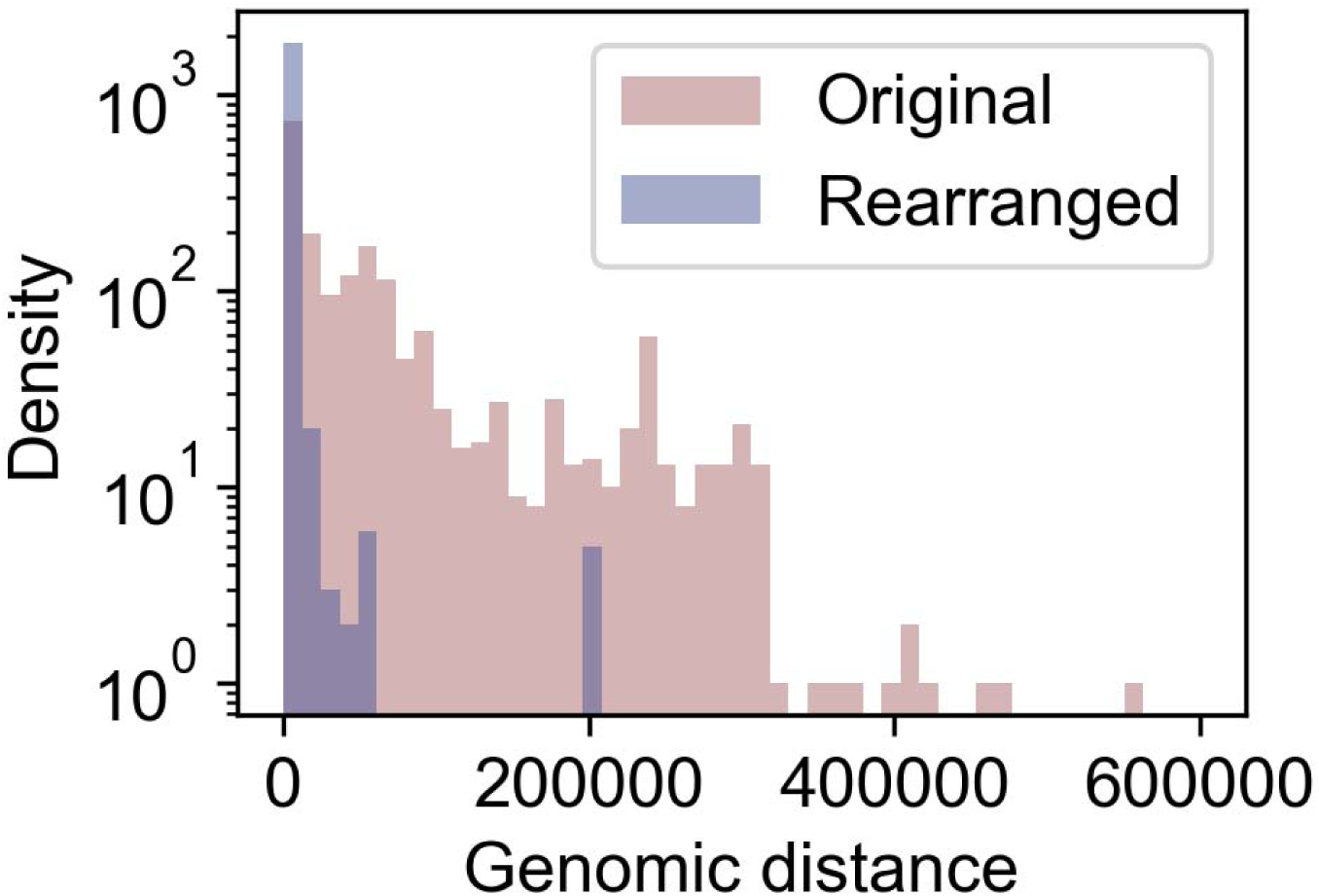
The distance between *opa* genes is higher in the original positions than after accounting for genomic rearrangements. The distances are calculated as the distance between each *opa* in the reference genome FA1090 to the *opa* that is closest in genomic position in all other isolates that had 11 *opa*.

**Figure S4:**
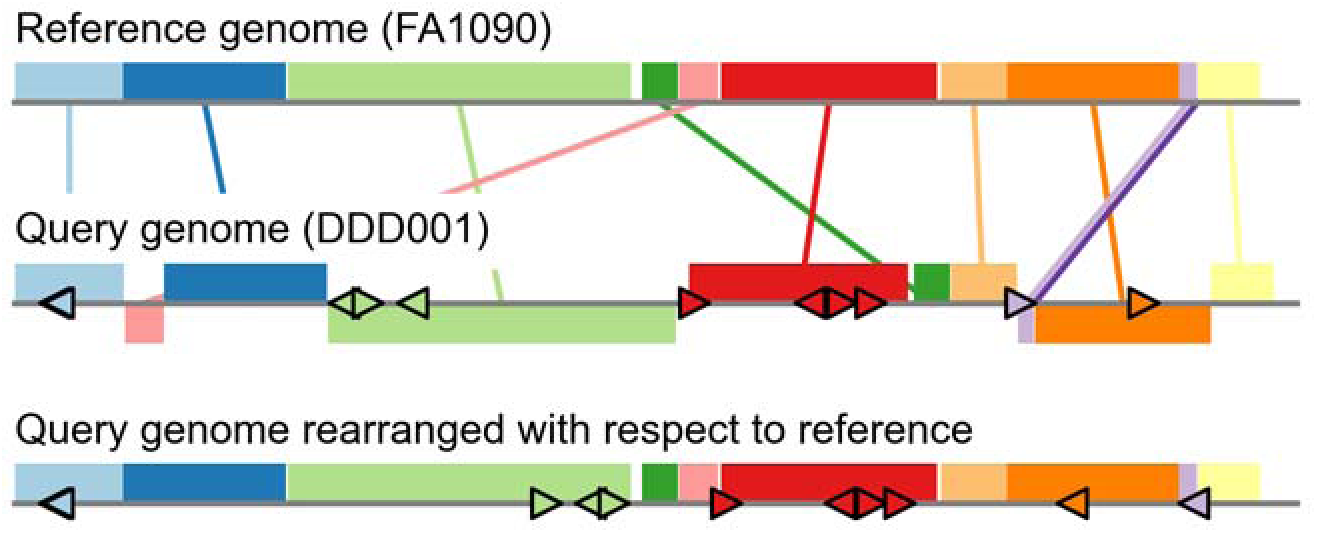
Schematic of genomic rearrangement procedure (see Methods). The shaded colored regions are the locally collinear blocks (LCBs). The LCBs that appear above the gray line are on the forward strand and those that appear below the gray line are on the reverse strand. A triangle pointing to the right indicates an *opa* gene on the forward strand and a triangle pointing to the left indicates an *opa* gene on the reverse strand.

**Figure S5:**
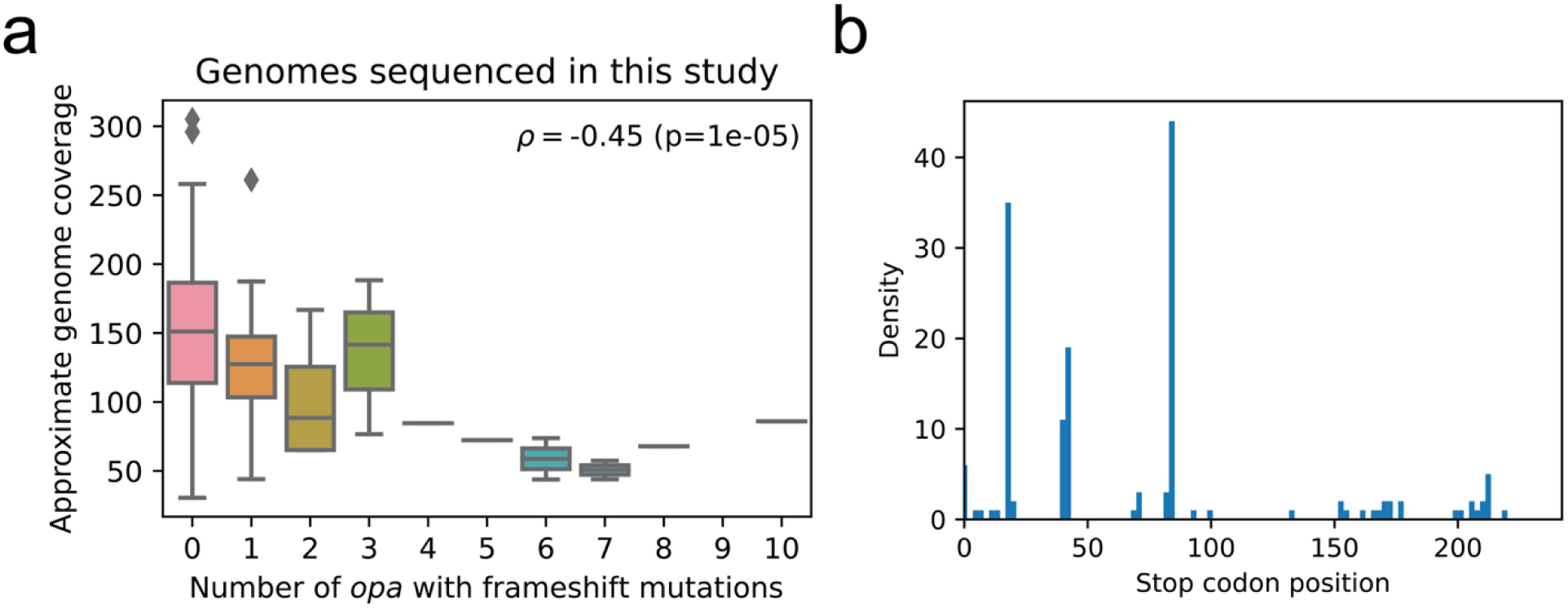
A subset of *opa* genes exhibit frameshift mutations after coding repeats leading to a premature stop codon. (a) The approximate genome sequencing coverage and number of *opa* in the genome with frameshift mutations downstream of the coding repeats. (b) The locations of the stop codons in the *opa* genes with frameshift mutations downstream of the coding repeats. The maximum value of the x-axis is set at the average length of *opa* amino acid sequences.

**Figure S6:**
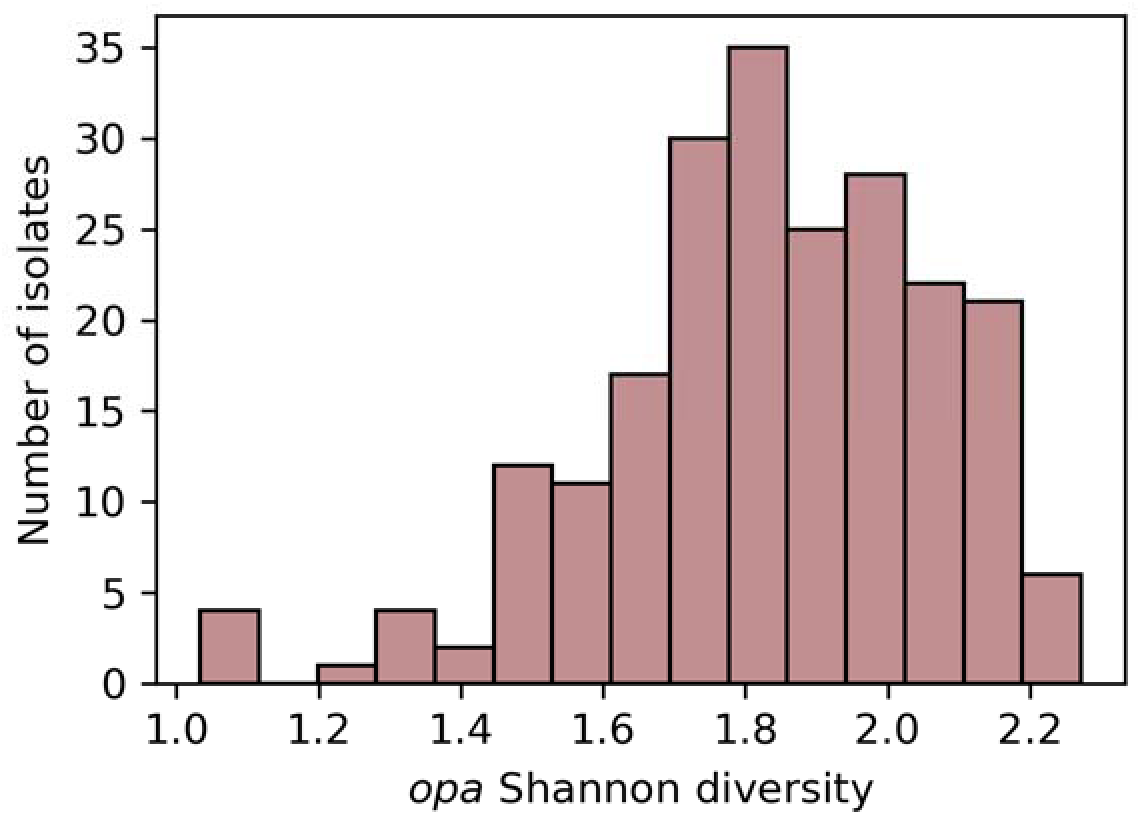
Shannon diversity of *opa* types within isolates.

**Figure S7:**
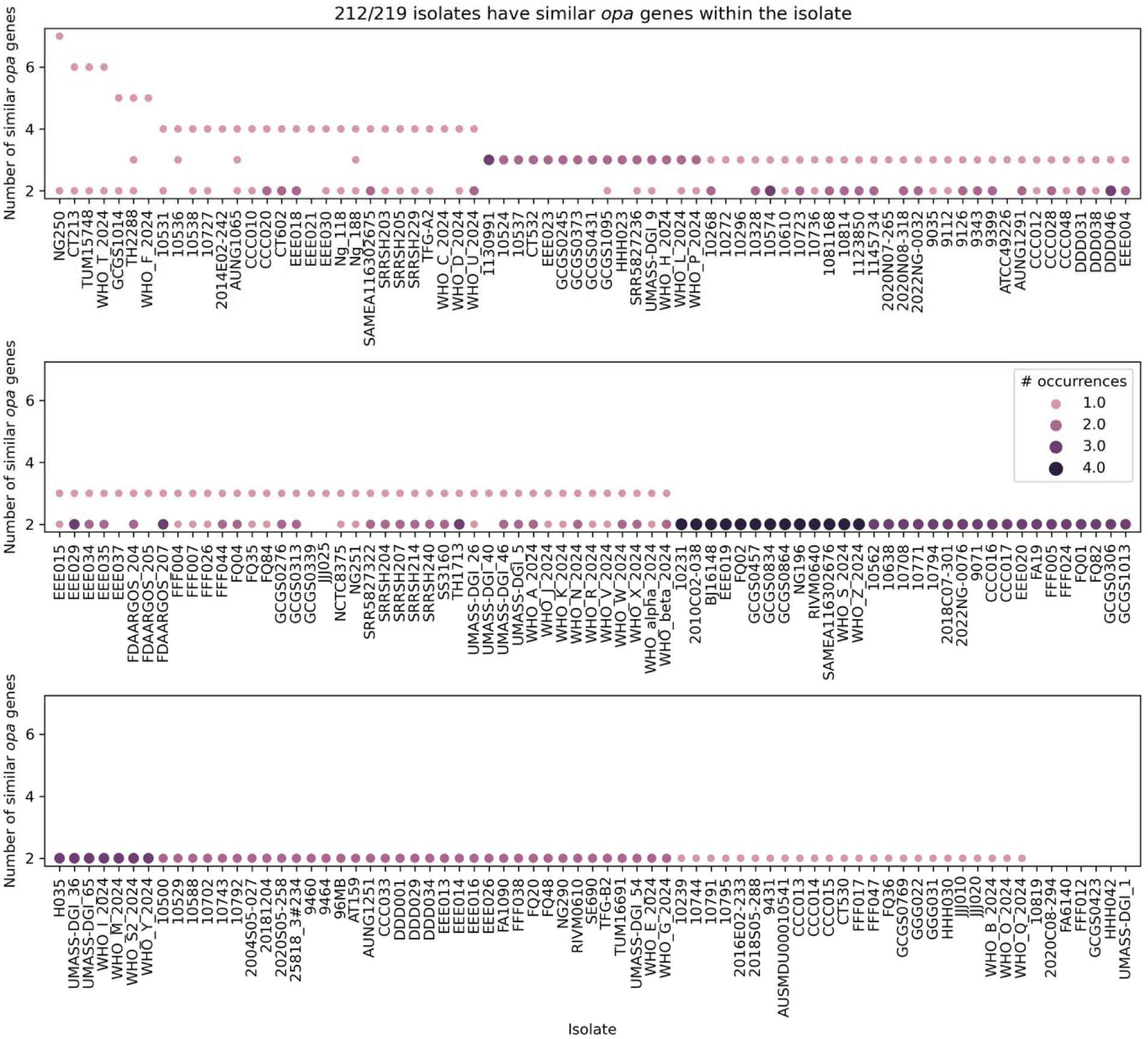
Genomes exhibit distinct patterns of similar *opa*s. All isolates with complete genomes are shown on the x-axis. The points indicate groups of *opa* genes in the same isolate with >95% amino acid sequence identity. The y-axis shows the number of similar *opa* genes in each group. The size and color of the point indicate the number of distinct groups of each size in the genome. The x-axis is sorted first by the maximum number of similar *opa* genes in any group and then by the maximum number of groups. The plot is split into three rows for readability.

**Figure S8:**
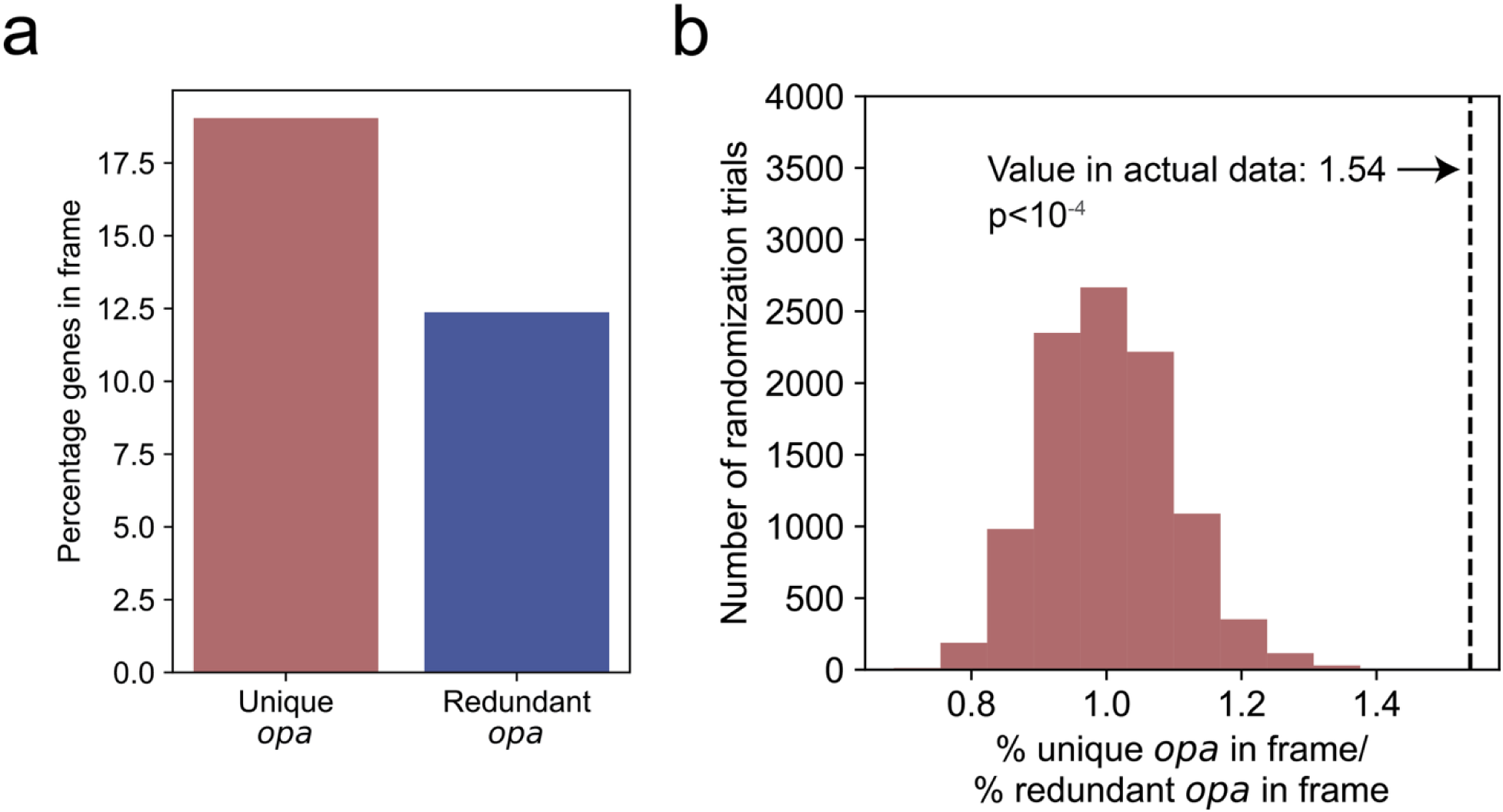
The *opa* that are unique in an isolate are more likely to be in frame than the *opa* that are redundant. (a) The percentage of *opa* that are in frame for *opa* that are unique within an isolate (<95% amino acid similarity) or redundant within an isolate (≥95% amino acid similarity). (b) The ratio of the percentage of unique *opa* in frame to the percentage of redundant *opa* in frame for 10^4^ randomizations of the data. The randomization procedure permuted which *opa* are labeled redundant, keeping the same total number of redundant *opa* across all isolates. Zero randomizations gave ratios as high as in the actual data (ratio of 1.54, indicated by the vertical dashed black line) yielding a p-value of less than 10^-4^.

**Figure S9.**
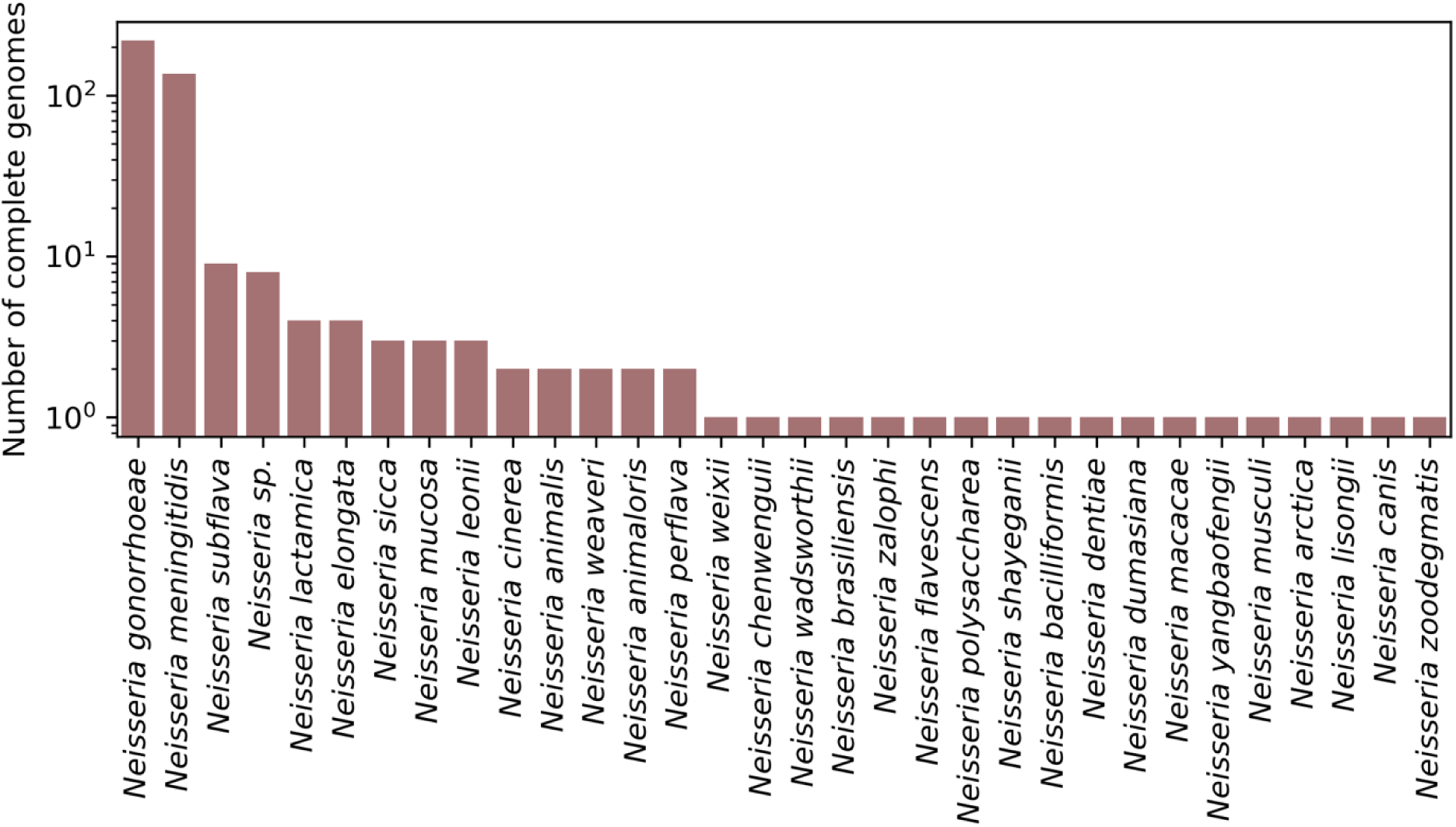
The number of publicly available complete genomes by species from the *Neisseria* genus.

**Figure S10:**
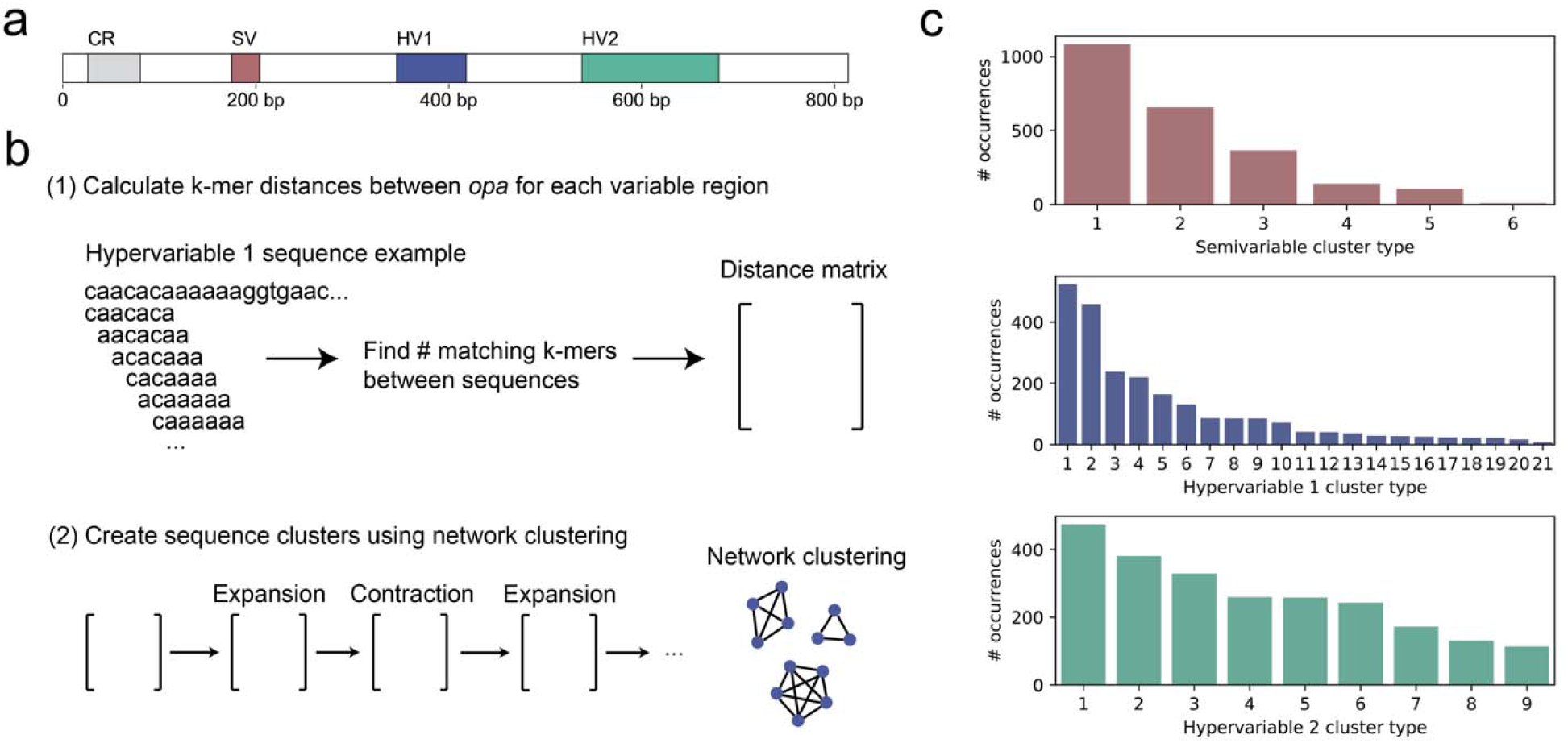
Network-based clustering approach for semivariable and hypervariable *opa* sequences. (a) Schematic of the *opa* gene. The exact length and locations of the gene features varies across *opa* genes; depicted here is FA1090 *opa1*. CR: coding repeat, SV: semivariable region, HV1: hypervariable 1 region, HV2: hypervariable 2 region. (b) Summary of the approach to clustering variable region sequences. For each variable region (semivariable, hypervariable 1, and hypervariable 2), we calculated the k-mer distances between all sequences using MASH, setting k such that the probability of finding a random k-mer in each sequence is 0.01. We performed successive rounds of inflation (expansion and contraction) on the distance matrix, which amplifies high values of the matrix and suppresses low values of the matrix. We chose the lowest inflation parameter that gave a stable clustering. (c) The distribution of the cluster types for the sequences in each variable region.

**Figure S11:**
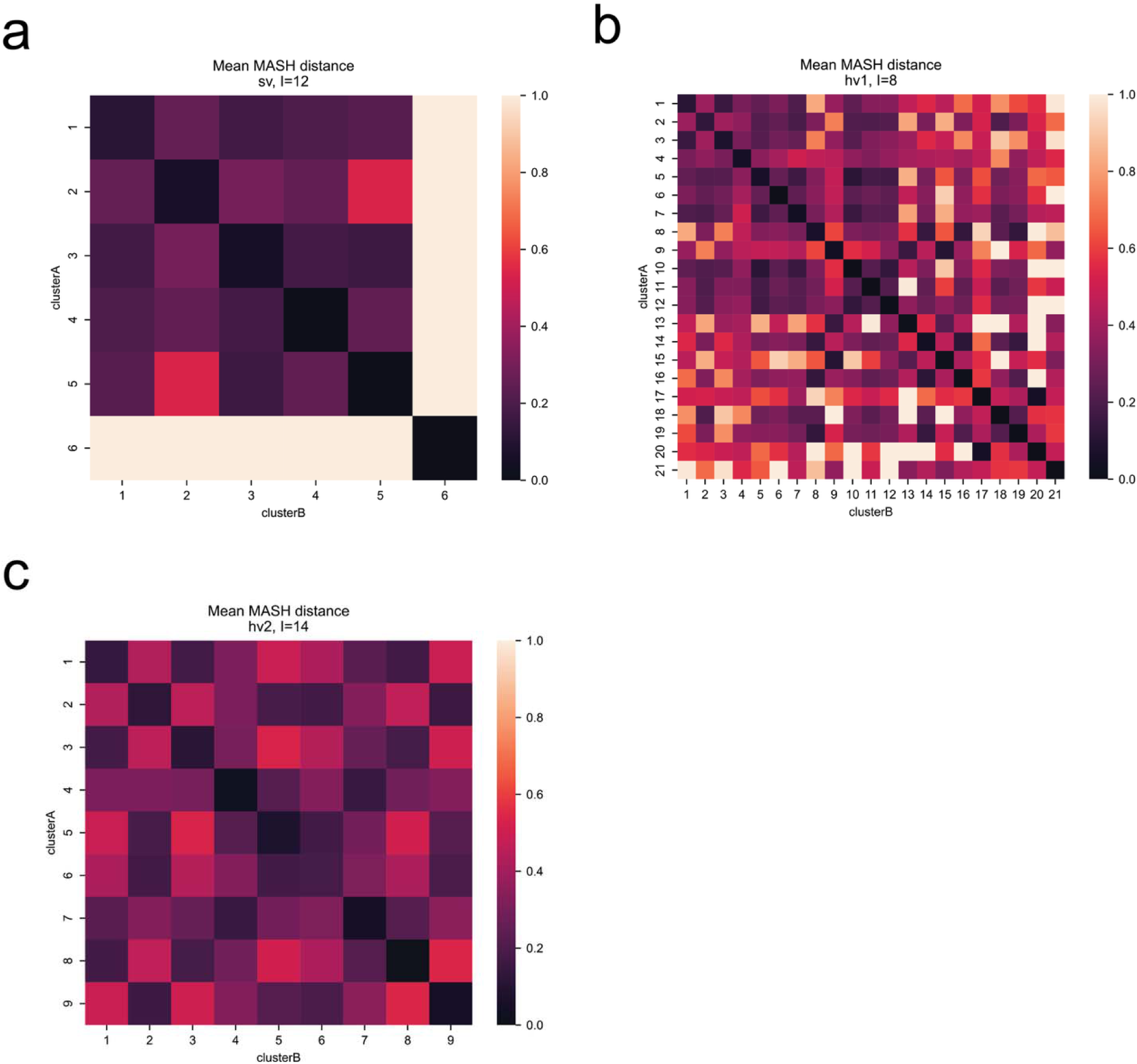
Sequences generally are more similar within clusters than between clusters and clusters have distinct sequence motifs. The mean nucleotide distance between pairs of sequences in the same and different clusters in the semivariable (a), hypervariable 1 (b), and hypervariable 2 (c) regions.

**Figure S12:**
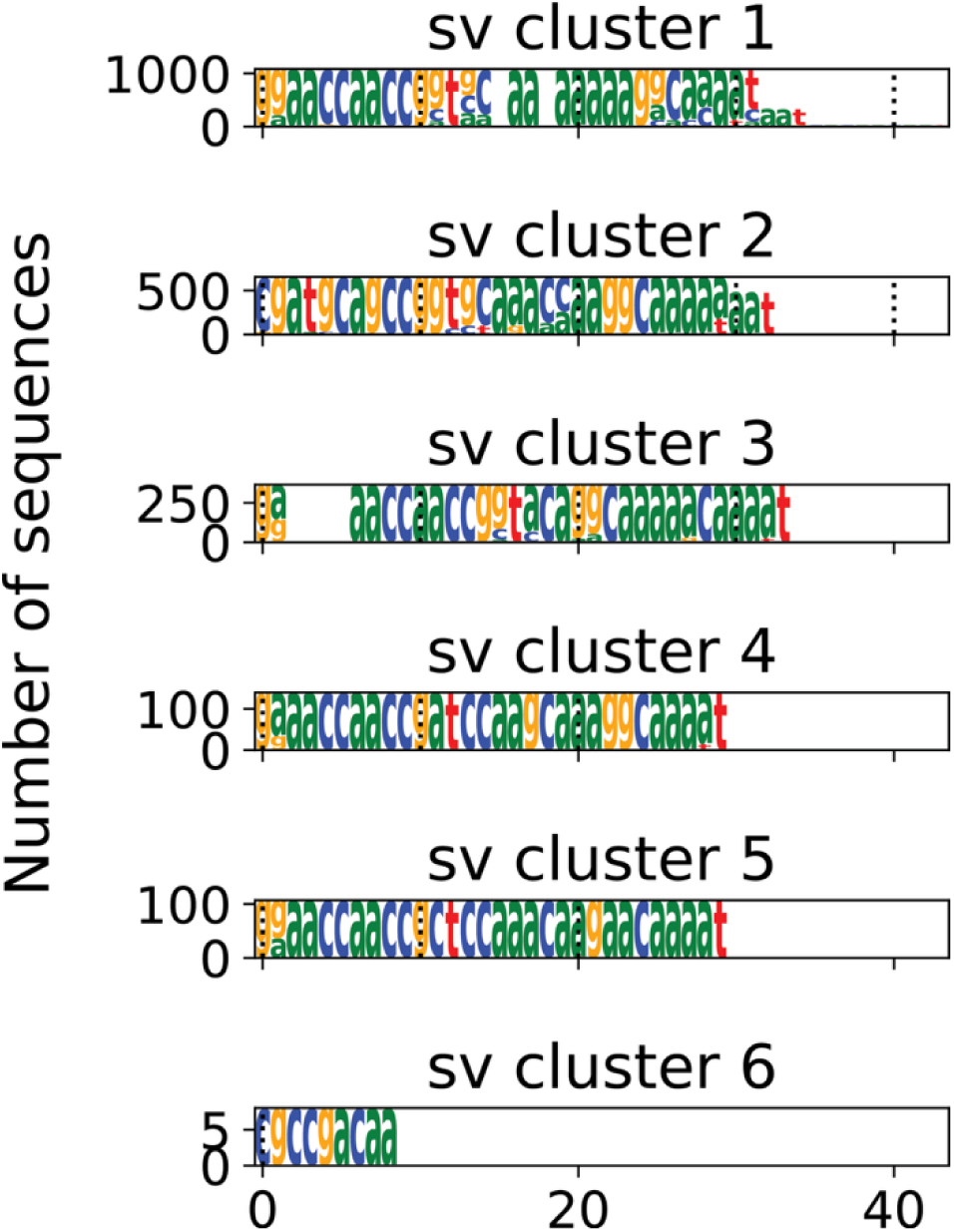
The sequence logos for each cluster of the semivariable sequences. The nucleotide sequences were aligned using MAFFT in each cluster. The height of the nucleotides represents the number of sequences with each nucleotide.

**Figure S13:**
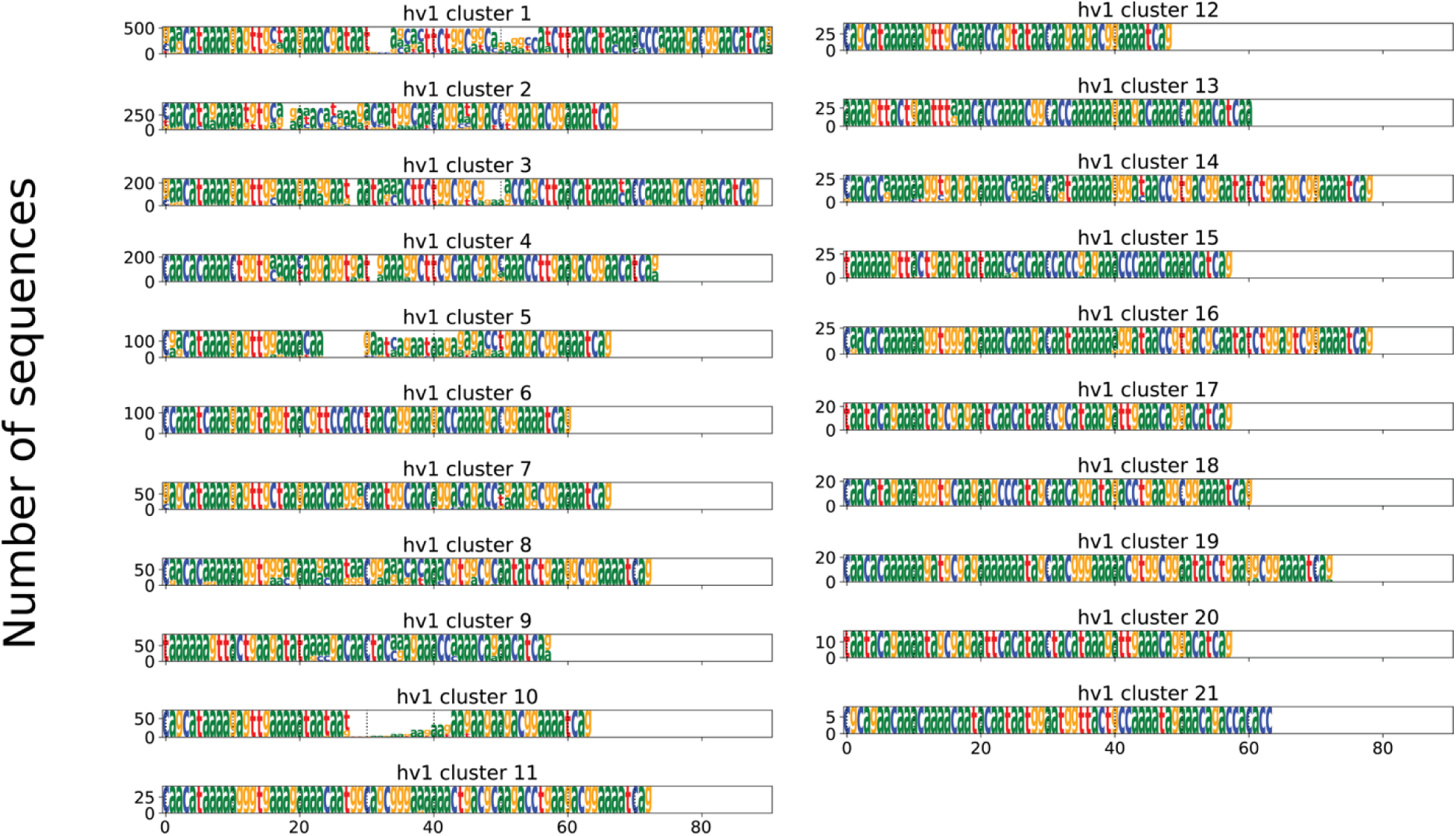
The sequence logos for each cluster of the hypervariable 1 sequences. The nucleotide sequences were aligned using MAFFT in each cluster. The height of the nucleotides represents the number of sequences with each nucleotide.

**Figure S14:**
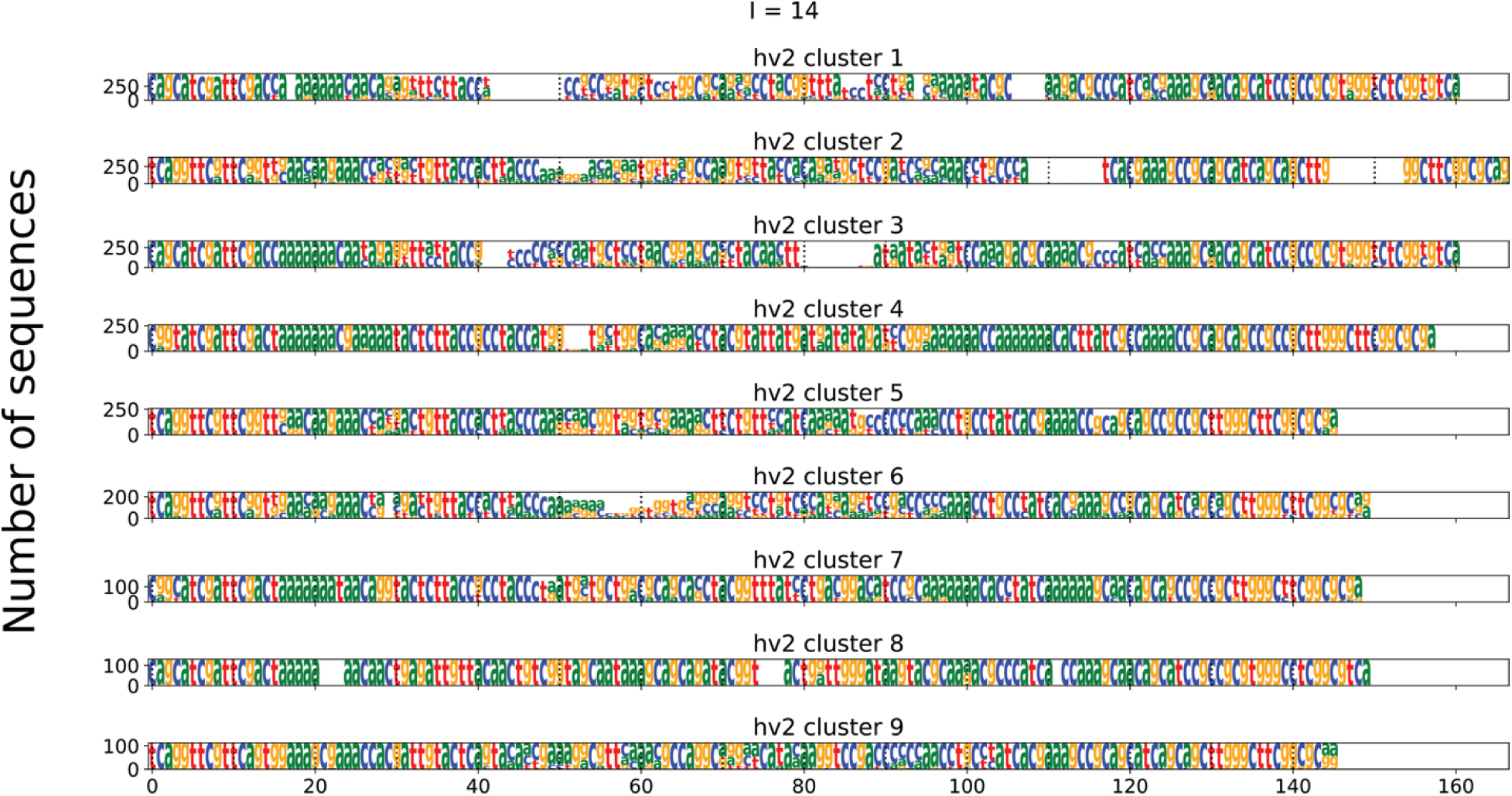
The sequence logos for each cluster of the hypervariable 2 sequences. The nucleotide sequences were aligned using MAFFT in each cluster. The height of the nucleotides represents the number of sequences with each nucleotide.

**Figure S15:**
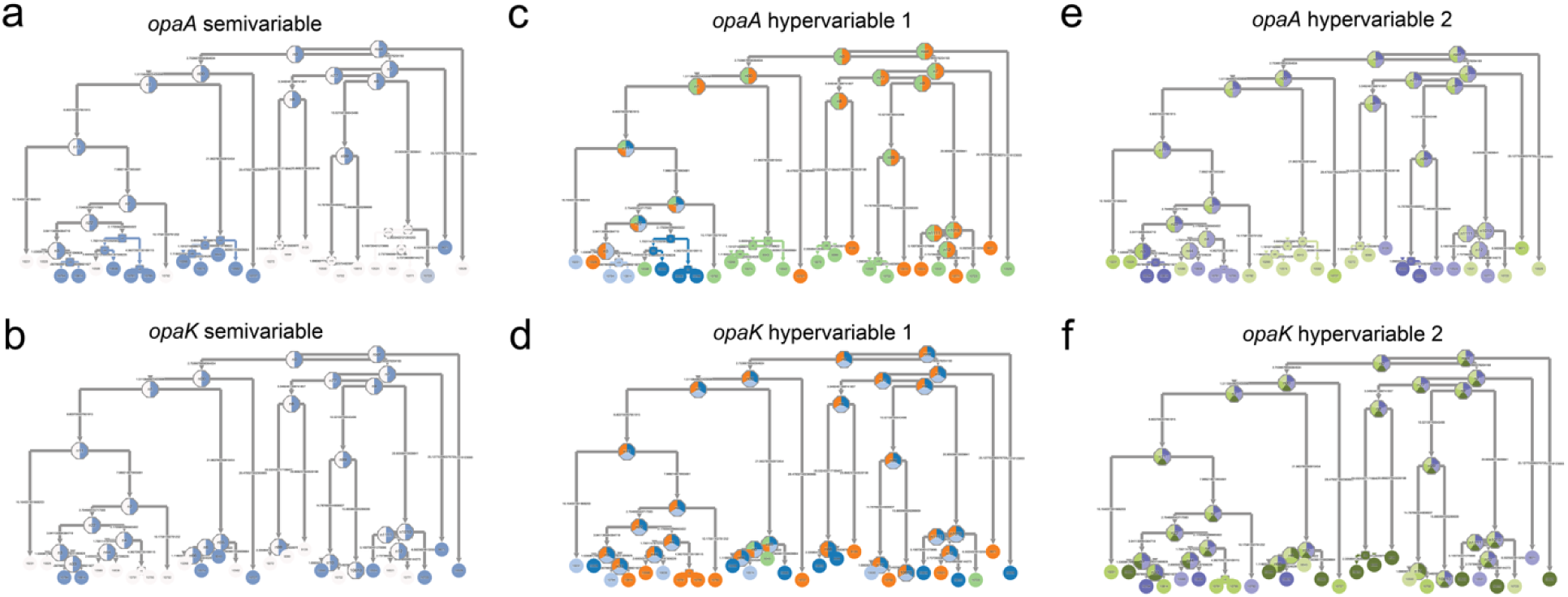
*opaA* and *opaK* have similar cluster types that change at different rates. The dated phylogeny of the subtree shown in Figure 6b annotated with the cluster types for the semivariable (a-b), hypervariable 1 (c-d), and hypervariable 2 (e-f) regions shown as colored circles for *opaA* (a, c, e) and *opaK* (b, d, f). The colors are comparable within each sequence region across loci (e.g., the same color scheme is used for the semivariable region of both *opaA* and *opaK*) but not between sequence regions. The colored circle annotations at the tips represent the cluster types of the isolates, and the colored circle annotations in the internal nodes represent the inferred ancestral state by PastML. The cluster types that are present in *opaA* and *opaK* are similar, but more closely related isolates have more similar cluster types in *opaA* compared to *opaK*.

**Figure S16:**
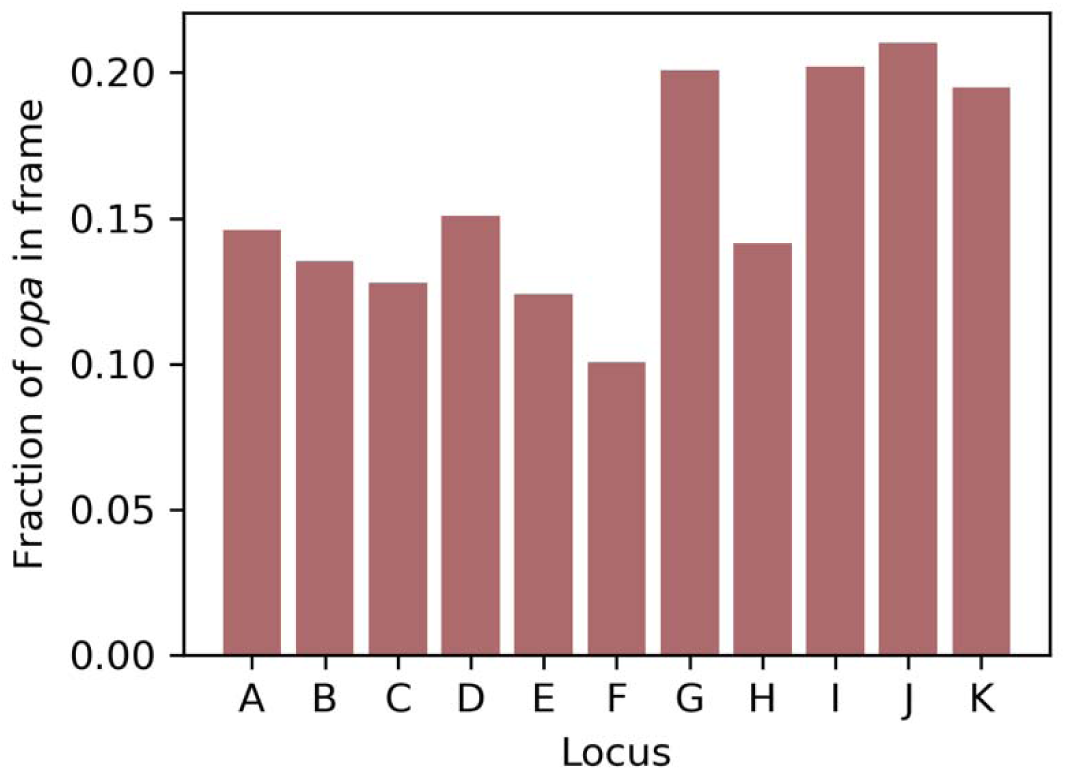
No significant differences in fraction in of *opa* in frame by locus. One-sided proportions *Z*-tests with a Bonferroni multiple hypothesis correction comparing the fraction of *opa* in frame at each locus (1) to the total fraction of *opa* in frame across all loci and (2) to the fraction of *opaK* alleles that are in frame are not significant.

**Figure S17:**
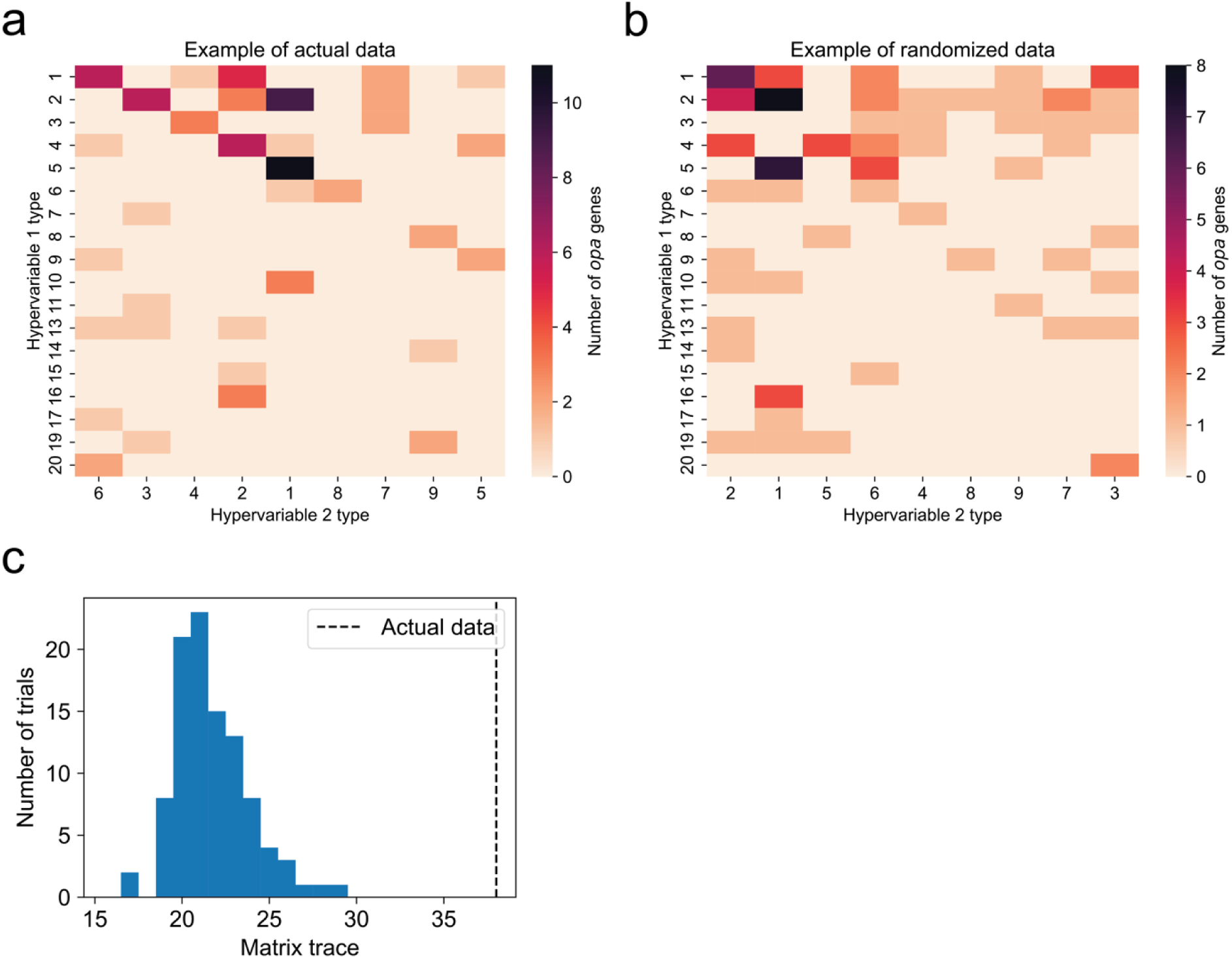
Non-random association of hypervariable 1 and hypervariable 2 allele types in *opa* genes, as shown for one representative random subset of 1 isolate per BAPS cluster. (a) The number of each combination of hypervariable 1 and hypervariable 2 types in *opa* alleles after accounting for isolate sampling and population structure. The columns of the matrix were rearranged to give the largest matrix trace (padding with columns of all zeros to make a square matrix). (b) The same data representation in (a) but after randomizing the assignment of hypervariable 2 types across the *opa* alleles. (c) The distribution of the maximum sum of the diagonal matrix elements (allowing for column rearrangements) in 100 randomizations of the hypervariable 2 type (blue histogram) compared to the actual data (black dashed line).

## Supplementary Information

### Supplementary Methods

#### Short-read sequencing

Short-read sequencing for EEE029 was performed in Bristow, Mortimer, et al. [68] and short-read sequencing of UMASS-DGI_65 was performed using the same method. Briefly, DNA was extracted from bacterial cells grown overnight on GCB-K plates at 37°C with 5% carbon dioxide using the Invitrogen PureLink Genomic DNA Mini Kit. Libraries were prepped and sequenced on the Illumina NextSeq 2000 sequencer at the Microbial Whole Genome Sequencing Center or the Bauer Core Facility at Harvard University.

#### Comparison of long-read assembly and polishing methods

To develop a pipeline for assembling complete genomes to minimize errors in the *opa* genes, we used the following assembly and polishing approaches using the filtered long-reads from Oxford Nanopore (see Methods) and short reads generated by Illumina NextSeq: Flye v2.9.5 [27], Medaka v2.0.1 (https://github.com/nanoporetech/medaka), Polypolish v0.6.0 [69], Pypolca v0.3.1 [69], and Autocycler v0.2.1 [25]. We tested the following workflows:

1. Long-read only assembly with Flye.
2. Long-read only assembly with Flye ➔ long-read polishing with Medaka
3. Long-read only assembly with Flye ➔ long-read polishing with Medaka ➔ short-read polishing with Polypolish default
4. Long-read only assembly with Flye ➔ long-read polishing with Medaka ➔ short-read polishing with Polypolish default followed by short-read polishing with Pypolca careful
5. Long-read only assembly with Autocycler
6. Long-read only assembly with Autocycler ➔ short-read polishing with Polypolish default
7. Long-read only assembly with Autocycler ➔ short-read polishing with Polypolish default ➔ short-read polishing with Pypolca careful

We used the genomes UMASS-DGI_65 and EEE029 which had sequencing depth of around 140x. To test how sequencing depth would affect the *opa* sequence accuracy, we randomly subsampled the reads using the seqtk package v1.4 (https://github.com/lh3/seqtk) to 15% (21x), 30% (42x), and 60% (84x), and compared to the results including 100% of the reads (140x). For each read subset, we assembled the genome using the approaches above and identified the *opa* sequences as described in the Methods. We aligned the resulting *opa* sequences using MAFFT v7.520 [35] with the default parameters.

#### More extensive testing of Autocycler with different read depth across diverse isolates

We selected 11 phylogenetically diverse isolates with at least 125x coverage using Treemmer v0.3 [70]. We created 10 random read subsets using the seqtk package at 50x, 75x, 100x, and 125x coverage. We assembled the genomes using the following approaches. We assembled the genomes (using long reads only) using Autocycler and followed the approach above to compare *opa* sequences.

#### Algorithm to search for opa genes in the complete genomes

We wrote a custom python script to identify *opa* genes. First, we searched for 3 to 100 tandem repeats of the CTCTT pentanucleotide with at most 2 substitutions across the entire sequence, which we will refer to as the coding repeats (CR), similar to the approach used in Bilek et al. [21]. We then searched for the unique conserved sequence near the stop codon (TGCGCTACCGCTTCTGAT) with at most 2 substitutions, which we will refer to as the term sequence. Pairs of CR and term sequences were matched if they were separated by less than 1200 bp in the genome. For term sequences that were unpaired, a more lenient search was performed for the upstream CR where we allowed up to 1 error for each CR unit (substitution, insertion, or deletion) to catch CR units that contained mutations. If there was still not a matching CR found, it could have been that the *opa* did not have a CR; in this case, we looked for the sequence found directly upstream of the CR consisting of a poly-A sequence of length 5-7 followed by CCTT and allowed one error (substitution, insertion, or deletion).

The start codon (ATG) was identified in the 50 bp upstream of the start of the CR sequence and the stop codon (TGA) was contained in the term sequence. Occasionally, the beginning and end of the CR region was not defined precisely with the above procedure due to errors in the CR (substitution, insertion, or deletion). Thus, the start of the CR was made more precise by finding the upstream sequence (see end of previous paragraph). If this sequence was not found, then the start of CR from earlier was used. Similarly, the end of the CR was made more precise by searching for the downstream sequence, which is CCG, allowing for the first C to be optional and 1 additional error.

#### Calculation of the randomness of hypervariable 1 and hypervariable 2 allele types

We dropped duplicate *opa* genes in the same genome as determined by having the same semivariable, hypervariable 1, and hypervariable 2 cluster type combinations to remove the effect of recent gene conversion events that have not yet had the time to be subject to selection.

We accounted for isolate sampling and *N. gonorrhoeae* population structure by partitioning the recombination-corrected phylogeny of complete genomes using fastbaps v1.0.8 [71] with the BAPS prior. We randomly selected 1 isolate from each BAPS cluster.

We determined the number of times each hypervariable 1 and hypervariable 2 cluster types appeared together in all *opa* genes in the subsampled isolates and created an association matrix where the rows were hypervariable 1 types, the columns were hypervariable 2 types, and the element values were the number of times each combination appeared. Because the number of hypervariable 1 and hypervariable 2 types were unequal, we padded columns with zeros to give a square matrix. We determined the degree of association by rearranging the columns of the association matrix to maximize the sum of diagonal entries using the Munkres algorithm (https://github.com/bmc/munkres?tab=readme-ov-file, v1.1.4). To determine whether the observed data could be random, we randomized the hypervariable 2 cluster types across the *opa* genes 100 times. We compared the maximum sum of diagonal elements from the observed data and the randomized data. We then repeated this process by performing 99 random subsamples of the isolates in each BAPS cluster.

#### Quantification of frameshift mutations downstream of the coding repeats leading to a premature stop codon

We identified all *opa* amino acid sequences that had a premature stop codon after the end of the coding repeat sequence. The approximate genome coverage was calculated as the number of input read bases divided by the number of consensus assembly bases. To identify the location of the frameshift mutations, the nucleotide sequences were aligned with MAFFT v7.520 [35] with the default parameters and visualized in Jalview v2.11.4.1 [36].

#### Comparison of long-read assembly and polishing methods

We first compared multiple assembly and polishing approaches, including Flye, Medaka, Polypolish, Pypolca, and Autocycler at 4 read depths (21x, 42x, 84x, and 140x) for two genomically diverse isolates. The *opa* sequences were identical across assemblies except for 1 SNP in UMASS-DGI_65 *opa7* and a 1 base insertion in EEE029 *opa11*, both of which occurred in methods 1 (long-read assembly with Flye) and 2 (long-read assembly with Flye followed by long-read polishing with Medaka) for a read depth of 21x (Figure S*1*). This analysis suggested Autocycler was the best performing assembly method and that polishing the Autocycler assemblies with short reads did not affect the *opa* sequences. However, because we only tested 2 genomes, we wanted to expand the analysis to include more isolates that are representative of *N. gonorrhoeae* diversity and to subset the reads to test lower read depths more systemically.

In our more extensive testing of Autocycler using 11 diverse isolates and 4 read depths (50x, 75x, 100x, and 125x) the *opa* sequences were identical across assemblies except for the following differences (always compared to 125x coverage assembly) (Figure S*2*):

- EEE023 *opa4* had 1 SNP in 10/10 assemblies with 50x coverage.
- FFF007 *opa11* was not detected in 10/10 assemblies with 50x coverage due to a missing stop codon and had multiple sequence differences in 10/10 assemblies with 75x coverage.
- GCGS0313 *opa1* had a 1 base deletion in 9/10 assemblies with 125x coverage
- GCGS0313 *opa5* had 1 SNP in 10/10 assemblies with 50x coverage

Despite these changes in a small number of *opa* sequences, we concluded that most *opa* sequences were identical across read depths and that Autocycler assemblies were good enough for our purposes of looking at diversity and evolution across a large set of *opa* genes.

## Supplementary Tables

**Table S1.** Publicly available complete genomes.

**Table S2.** Complete genomes sequenced in this study.

**Table S3.** References for publicly available *N. gonorrhoeae* short-read sequencing data meeting quality control thresholds for selection of representative draft genomes.

**Table S4.** Representative *N. gonorrhoeae* draft genomes.

**Table S5.** Publicly available *Neisseria* species complete genomes.

## References

1. WHO bacterial priority pathogens list, 2024: Bacterial pathogens of public health importance to guide research, development and strategies to prevent and control antimicrobial resistance. World Health Organization; 17 May 2024 [cited 12 Aug 2025]. Available: https://www.who.int/publications/i/item/9789240093461

2. Unemo M, Seifert HS, Hook EW 3rd, Hawkes S, Ndowa F, Dillon J-AR. Gonorrhoea. Nat Rev Dis Primers. 2019;5: 79.

3. Schmidt KA, Schneider H, Lindstrom JA, Boslego JW, Warren RA, Van de Verg L, et al. Experimental gonococcal urethritis and reinfection with homologous gonococci in male volunteers. Sex Transm Dis. 2001;28: 555–564.

4. Belcher T, Rollier CS, Dold C, Ross JDC, MacLennan CA. Immune responses to Neisseria gonorrhoeae and implications for vaccine development. Front Immunol. 2023;14: 1248613.

5. Fox KK, Thomas JC, Weiner DH, Davis RH, Sparling PF, Cohen MS. Longitudinal evaluation of serovar-specific immunity to Neisseria gonorrhoeae. Am J Epidemiol. 1999;149: 353–358.

6. de Korne-Elenbaas J, Bruisten SM, de Vries HJC, van Dam AP. Within-Host Genetic Variation in Neisseria gonorrhoeae over the Course of Infection. Microbiol Spectr. 2022;10: e0031322.

7. Quillin SJ, Seifert HS. Neisseria gonorrhoeae host adaptation and pathogenesis. Nat Rev Microbiol. 2018;16: 226–240.

8. Black WJ, Schwalbe RS, Nachamkin I, Cannon JG. Characterization of Neisseria gonorrhoeae protein II phase variation by use of monoclonal antibodies. Infect Immun. 1984;45: 453–457.

9. Malorny B, Morelli G, Kusecek B, Kolberg J, Achtman M. Sequence diversity, predicted two-dimensional protein structure, and epitope mapping of neisserial Opa proteins. J Bacteriol. 1998;180: 1323–1330.

10. Dempsey JA, Litaker W, Madhure A, Snodgrass TL, Cannon JG. Physical map of the chromosome of Neisseria gonorrhoeae FA1090 with locations of genetic markers, including opa and pil genes. J Bacteriol. 1991;173: 5476–5486.

11. Connell TD, Black WJ, Kawula TH, Barritt DS, Dempsey JA, Kverneland K Jr, et al. Recombination among protein II genes of Neisseria gonorrhoeae generates new coding sequences and increases structural variability in the protein II family. Mol Microbiol. 1988;2: 227–236.

12. Chen T, Belland RJ, Wilson J, Swanson J. Adherence of pilus-Opa+ gonococci to epithelial cells in vitro involves heparan sulfate. J Exp Med. 1995;182: 511–517.

13. van Putten JP, Paul SM. Binding of syndecan-like cell surface proteoglycan receptors is required for Neisseria gonorrhoeae entry into human mucosal cells. EMBO J. 1995;14: 2144–2154.

14. Virji M, Evans D, Hadfield A, Grunert F, Teixeira AM, Watt SM. Critical determinants of host receptor targeting by Neisseria meningitidis and Neisseria gonorrhoeae: identification of Opa adhesiotopes on the N-domain of CD66 molecules. Mol Microbiol. 1999;34: 538–551.

15. Sadarangani M, Pollard AJ, Gray-Owen SD. Opa proteins and CEACAMs: pathways of immune engagement for pathogenic Neisseria. FEMS Microbiol Rev. 2011;35: 498–514.

16. Gómez-Duarte OG, Dehio M, Guzmán CA, Chhatwal GS, Dehio C, Meyer TF. Binding of vitronectin to opa-expressing Neisseria gonorrhoeae mediates invasion of HeLa cells. Infect Immun. 1997;65: 3857–3866.

17. Duensing TD, van Putten JP. Vitronectin mediates internalization of Neisseria gonorrhoeae by Chinese hamster ovary cells. Infect Immun. 1997;65: 964–970.

18. Pantelic M, Kim Y-J, Bolland S, Chen I, Shively J, Chen T. Neisseria gonorrhoeae kills carcinoembryonic antigen-related cellular adhesion molecule 1 (CD66a)-expressing human B cells and inhibits antibody production. Infect Immun. 2005;73: 4171–4179.

19. Bhat KS, Gibbs CP, Barrera O, Morrison SG, Jähnig F, Stern A, et al. The opacity proteins of Neisseria gonorrhoeae strain MS11 are encoded by a family of 11 complete genes. Mol Microbiol. 1991;5: 1889–1901.

20. Stern A, Brown M, Nickel P, Meyer TF. Opacity genes in Neisseria gonorrhoeae: control of phase and antigenic variation. Cell. 1986;47: 61–71.

21. Bilek N, Ison CA, Spratt BG. Relative contributions of recombination and mutation to the diversification of the opa gene repertoire of Neisseria gonorrhoeae. J Bacteriol. 2009;191: 1878–1890.

22. van der Ley P. Three copies of a single protein II-encoding sequence in the genome of Neisseria gonorrhoeae JS3: evidence for gene conversion and gene duplication. Mol Microbiol. 1988;2: 797–806.

23. Kellogg DS Jr, Peacock WL Jr, Deacon WE, Brown L, Pirkle CI. Neisseria gonorrhoeae i. J Bacteriol. 1963;85: 1274–1279.

24. Oxford Nanopore Technologies. Dorado. 2024. Available: https://github.com/nanoporetech/dorado

25. Wick RR, Howden BP, Stinear TP. Autocycler: long-read consensus assembly for bacterial genomes. bioRxiv. 2025. p. 2025.05.12.653612. doi:10.1101/2025.05.12.653612

26. Koren S, Walenz BP, Berlin K, Miller JR, Bergman NH, Phillippy AM. Canu: scalable and accurate long-read assembly via adaptive k-mer weighting and repeat separation. Genome Res. 2017;27: 722–736.

27. Kolmogorov M, Yuan J, Lin Y, Pevzner PA. Assembly of long, error-prone reads using repeat graphs. Nat Biotechnol. 2019;37: 540–546.

28. Li H. Minimap and miniasm: fast mapping and de novo assembly for noisy long sequences. Bioinformatics. 2016;32: 2103–2110.

29. Chen Y, Nie F, Xie S-Q, Zheng Y-F, Dai Q, Bray T, et al. Efficient assembly of nanopore reads via highly accurate and intact error correction. Nat Commun. 2021;12: 60.

30. Hu J, Wang Z, Sun Z, Hu B, Ayoola AO, Liang F, et al. NextDenovo: an efficient error correction and accurate assembly tool for noisy long reads. Genome Biol. 2024;25: 107.

31. Vaser R, Šikić M. Time- and memory-efficient genome assembly with Raven. Nat Comput Sci. 2021;1: 332–336.

32. Durbin R, De Sanctis B, Blumer M. Rotate: A command-line program to rotate circular DNA sequences to start at a given position or string. Wellcome Open Res. 2023;8: 401.

33. Seemann T. Prokka: rapid prokaryotic genome annotation. Bioinformatics. 2014;30: 2068–2069.

34. Page AJ, Cummins CA, Hunt M, Wong VK, Reuter S, Holden MTG, et al. Roary: rapid large-scale prokaryote pan genome analysis. Bioinformatics. 2015;31: 3691–3693.

35. Katoh K, Standley DM. MAFFT multiple sequence alignment software version 7: improvements in performance and usability. Mol Biol Evol. 2013;30: 772–780.

36. Waterhouse AM, Procter JB, Martin DMA, Clamp M, Barton GJ. Jalview Version 2--a multiple sequence alignment editor and analysis workbench. Bioinformatics. 2009;25: 1189–1191.

37. Darling AE, Mau B, Perna NT. progressiveMauve: multiple genome alignment with gene gain, loss and rearrangement. PLoS One. 2010;5: e11147.

38. Huang W, Li L, Myers JR, Marth GT. ART: a next-generation sequencing read simulator. Bioinformatics. 2012;28: 593–594.

39. Li H. Aligning sequence reads, clone sequences and assembly contigs with BWA-MEM. arXiv [q-bio.GN]. 2013. Available: http://arxiv.org/abs/1303.3997

40. Danecek P, Bonfield JK, Liddle J, Marshall J, Ohan V, Pollard MO, et al. Twelve years of SAMtools and BCFtools. Gigascience. 2021;10. doi:10.1093/gigascience/giab008

41. Okonechnikov K, Conesa A, García-Alcalde F. Qualimap 2: advanced multi-sample quality control for high-throughput sequencing data. Bioinformatics. 2016;32: 292–294.

42. Walker BJ, Abeel T, Shea T, Priest M, Abouelliel A, Sakthikumar S, et al. Pilon: an integrated tool for comprehensive microbial variant detection and genome assembly improvement. PLoS One. 2014;9: e112963.

43. Lees JA, Harris SR, Tonkin-Hill G, Gladstone RA, Lo SW, Weiser JN, et al. Fast and flexible bacterial genomic epidemiology with PopPUNK. Genome Res. 2019;29: 304–316.

44. Croucher NJ, Page AJ, Connor TR, Delaney AJ, Keane JA, Bentley SD, et al. Rapid phylogenetic analysis of large samples of recombinant bacterial whole genome sequences using Gubbins. Nucleic Acids Res. 2015;43: e15.

45. Kozlov AM, Darriba D, Flouri T, Morel B, Stamatakis A. RAxML-NG: a fast, scalable and user-friendly tool for maximum likelihood phylogenetic inference. Bioinformatics. 2019;35: 4453–4455.

46. Ondov BD, Treangen TJ, Melsted P, Mallonee AB, Bergman NH, Koren S, et al. Mash: fast genome and metagenome distance estimation using MinHash. Genome Biol. 2016;17: 132.

47. Simonsen M, Mailund T, Pedersen CNS. Rapid neighbour-joining. Lecture Notes in Computer Science. Berlin, Heidelberg: Springer Berlin Heidelberg; 2008. pp. 113–122.

48. Van Dongen S. Graph clustering via a discrete uncoupling process. SIAM J Matrix Anal Appl. 2008;30: 121–141.

49. Enright AJ, Van Dongen S, Ouzounis CA. An efficient algorithm for large-scale detection of protein families. Nucleic Acids Res. 2002;30: 1575–1584.

50. Tareen A, Kinney JB. Logomaker: beautiful sequence logos in Python. Bioinformatics. 2020;36: 2272–2274.

51. Bouckaert R, Vaughan TG, Barido-Sottani J, Duchêne S, Fourment M, Gavryushkina A, et al. BEAST 2.5: An advanced software platform for Bayesian evolutionary analysis. PLoS Comput Biol. 2019;15: e1006650.

52. Ishikawa SA, Zhukova A, Iwasaki W, Gascuel O. A fast likelihood method to reconstruct and visualize ancestral scenarios. Mol Biol Evol. 2019;36: 2069–2085.

53. Letunic I, Bork P. Interactive Tree of Life (iTOL) v6: recent updates to the phylogenetic tree display and annotation tool. Nucleic Acids Res. 2024;52: W78–W82.

54. Köster J, Rahmann S. Snakemake--a scalable bioinformatics workflow engine. Bioinformatics. 2012;28: 2520–2522.

55. Unemo M, Sánchez-Busó L, Golparian D, Jacobsson S, Shimuta K, Lan PT, et al. The novel 2024 WHO Neisseria gonorrhoeae reference strains for global quality assurance of laboratory investigations and superseded WHO N. gonorrhoeae reference strains-phenotypic, genetic and reference genome characterization. J Antimicrob Chemother. 2024. doi:10.1093/jac/dkae176

56. Wadsworth CB, Arnold BJ, Sater MRA, Grad YH. Azithromycin Resistance through Interspecific Acquisition of an Epistasis-Dependent Efflux Pump Component and Transcriptional Regulator in Neisseria gonorrhoeae. MBio. 2018;9. doi:10.1128/mBio.01419-18

57. Unitt A, Maiden M, Harrison O. Characterizing the diversity and commensal origins of penA mosaicism in the genus Neisseria. Microb Genom. 2024;10. doi:10.1099/mgen.0.001209

58. Grad YH, Kirkcaldy RD, Trees D, Dordel J, Harris SR, Goldstein E, et al. Genomic epidemiology of Neisseria gonorrhoeae with reduced susceptibility to cefixime in the USA: a retrospective observational study. Lancet Infect Dis. 2014;14: 220–226.

59. Belland RJ, Morrison SG, Carlson JH, Hogan DM. Promoter strength influences phase variation of neisserial opa genes. Mol Microbiol. 1997;23: 123–135.

60. James JF, Swanson J. Studies on gonococcus infection. XIII. Occurrence of color/opacity colonial variants in clinical cultures. Infect Immun. 1978;19: 332–340.

61. James JF, Swanson J. Color/opacity colonial variants of Neisseria gonorrhoeae and their relationship to the menstrual cycle. In: G. F. Brooks, E. C. Gotschlich, K. K. Holmes, W. D. Sawyer, and F. E. Young, editor. Immunobiology of Neisseria gonorrhoeae. Annals of Internal Medicine; 1978. pp. 338–343.

62. Salit IE. The differential susceptibility of gonococcal opacity variants to sex hormones. Can J Microbiol. 1982;28: 301–306.

63. Jerse AE, Cohen MS, Drown PM, Whicker LG, Isbey SF, Seifert HS, et al. Multiple gonococcal opacity proteins are expressed during experimental urethral infection in the male. J Exp Med. 1994;179: 911–920.

64. Swanson J, Barrera O, Sola J, Boslego J. Expression of outer membrane protein II by gonococci in experimental gonorrhea. J Exp Med. 1988;168: 2121–2129.

65. Callaghan MJ, Jolley KA, Maiden MCJ. Opacity-associated adhesin repertoire in hyperinvasive Neisseria meningitidis. Infect Immun. 2006;74: 5085–5094.

66. Callaghan MJ, Buckee CO, Jolley KA, Kriz P, Maiden MCJ, Gupta S. The effect of immune selection on the structure of the meningococcal opa protein repertoire. PLoS Pathog. 2008;4: e1000020.

67. Bos MP, Kao D, Hogan DM, Grant CCR, Belland RJ. Carcinoembryonic antigen family receptor recognition by gonococcal Opa proteins requires distinct combinations of hypervariable Opa protein domains. Infect Immun. 2002;70: 1715–1723.

68. Bristow CC, Mortimer TD, Morris S, Grad YH, Soge OO, Wakatake E, et al. Whole-genome sequencing to predict antimicrobial susceptibility profiles in Neisseria gonorrhoeae. J Infect Dis. 2023;227: 917–925.

69. Bouras G, Judd LM, Edwards RA, Vreugde S, Stinear TP, Wick RR. How low can you go? Short-read polishing of Oxford Nanopore bacterial genome assemblies. bioRxiv. 2024. p. 2024.03.07.584013. doi:10.1101/2024.03.07.584013

70. Menardo F, Loiseau C, Brites D, Coscolla M, Gygli SM, Rutaihwa LK, et al. Treemmer: a tool to reduce large phylogenetic datasets with minimal loss of diversity. BMC Bioinformatics. 2018;19: 164.

71. Tonkin-Hill G, Lees JA, Bentley SD, Frost SDW, Corander J. Fast hierarchical Bayesian analysis of population structure. Nucleic Acids Res. 2019;47: 5539–5549.

